# Rescuing epileptic and behavioral alterations in a Dravet syndrome mouse model by inhibiting eukaryotic eEF2K

**DOI:** 10.1101/2021.07.07.451562

**Authors:** Stefania Beretta, Luisa Ponzoni, Laura Gritti, Paolo Scalmani, Massimo Mantegazza, Mariaelvina Sala, Chiara Verpelli, Carlo Sala

**Affiliations:** CNR Neuroscience Institute, Milan, 20854 Vedano al Lambro, Italy; L’Unità Operativa Complessa di Epilettologia Clinica e Sperimentale, Foundation Istituto di Ricerca e Cura a Carattere Scientifico (IRCCS), Neurological Institute Carlo Besta, 20133 Milan, Italy; CNRS UMR 7275, Institut National de la Santé et de la Recherche Médicale, LabEx ICST, Institute of Molecular and Cellular Pharmacology (IPMC), Université Côte d’Azur (UCA), 06560 Valbonne-Sophia Antipolis, France

**Keywords:** inhibitory synapses, protein translation, EEG, *SCN1A* gene

## Abstract

Dravet Syndrome is a severe childhood pharmacoresistant epileptic disorder caused mainly by mutations in the *SCN1A* gene, which encodes for the α1 subunit of the type I voltage-gated sodium channel (Na_V_1.1), that cause imbalance between excitation and inhibition in the brain. We recently found that eEF2K knock out mice displayed enhanced GABAergic transmission and tonic inhibition and were less susceptible to epileptic seizures. In Scn1a+/- mice, a mouse model of the Dravet syndrome, we found that the activity of eEF2K/eEF2 pathway was enhanced. Then, we demonstrated that both genetic deletion and pharmacological inhibition of eEF2K were able to reduce the epileptic phenotype of Scn1a+/- mice. Interestingly we also found that motor coordination defect, memory impairments, and stereotyped behavior of the *Scn1a*^+/-^ mice were reverted by eEF2K deletion. The analysis of spontaneous inhibitory postsynaptic currents (sIPSCs) suggested that the rescue of the pathological phenotype was driven by the potentiation of GABAergic synapses. Our data indicate that pharmacological inhibition of eEF2K could represent a novel therapeutic intervention for treating epilepsy and related comorbidities in the Dravet syndrome.

## Introduction

Dravet syndrome (DS), also known as Severe Myoclonic Epilepsy in Infancy (SMEI), is a severe childhood disorder that typically presents in the first years of life with pharmacoresistant epilepsy followed by devastating effects on cognitive development. DS typically presents with febrile seizures that later develop to severe partial or generalized tonic-clonic seizures, myoclonic seizures, atypical absences, and focal seizures, as well as episodes of status epilepticus (Dravet, 2011a, b; Bender et al., 2012). Starting from the second year of life patients develop important comorbidities such as cognitive impairment, including intellectual disability and autistic traits (Genton et al., 2011; Ragona, 2011; Bender et al., 2016), behavioral disturbances, including psychomotor delay, ataxia, sleep disorder, impairment in visuospatial and language development (Dravet et al., 2005; Jansen et al., 2006), and they also have higher incidence of premature death. Approximately 80% of patients with Dravet syndrome carry mutations in the *SCN1A* gene which encodes for the α1 subunit of the type I voltage-gated sodium channel (Nav1.1) (Claes et al., 2001; Mantegazza et al., 2005). Mutations associated with Dravet syndrome are randomly distributed along the gene and include missense (40%), nonsense/truncated (40%) and the remaining 20% are frameshift mutations, which can be the result of insertion, duplication and deletion of nucleotide (Marini et al., 2011; Bender et al., 2012; Catterall, 2014). Na_V_1.1 containing channels, despite being broadly expressed in different neurons of the brain including pyramidal neurons, are highly expressed in GABAergic interneurons in hippocampus and cerebral cortex. Loss-of-function of Na_V_1.1, reducing sodium current, greatly impairs the ability of these interneurons to fire action potentials at high frequency and therefore reduces their phasic release of GABA (Yu et al., 2006; Ogiwara et al., 2007; Bender et al., 2012; Mantegazza et al., 2021). This results in an imbalance between excitation and inhibition that leads to hyperexcitability and seizures (Bender et al., 2012) and suggests that a promising therapy for DS might be based on normalizing this excitation/inhibition unbalance.

eEF2K, previously known as calcium/calmodulin-dependent protein Kinase III (CaMKIII), is a ubiquitous protein kinase involved in the control of mRNA translation, whose catalytic activity is Ca^2+^-dependent. Upon activation, eEF2K phosphorylates and inhibits eukaryotic elongation factor 2 (eEF2), leading to inhibition of mRNA translation at the level of elongation (Ryazanov et al., 1988; Browne and Proud, 2002). We recently demonstrated that eEF2K deletion in mice caused an enhancement in GABAergic transmission that was accompanied by an increased resistance to epilepsy (Heise et al., 2017). Accordingly, we showed that eEF2K deletion, in a mouse model of human epilepsy, the Synapsin1 knock out mice rescued the epileptic phenotype (Heise et al., 2017). In this study we investigated the effect of inhibition of eEF2K on the epileptic and behavioral phenotype of Scn1a+/- mice, a murine model of DS (Yu et al., 2006; Han et al., 2012). Our results demonstrate that genetic deletion of eEF2K in Scn1a+/- mice rescued both electroencephalography (EEG) and behavioral alterations. Additionally, we show that pharmacological inhibition of eEF2K ameliorated the altered EEG phenotype, demonstrating that treatments aimed to decrease eEF2K activity might represent a new approach for treating patients that are affected by DS.

## Results

### Higher level of phosphorylate eEF2 (P-eEF2) in cortex and hippocampus of Scn1a+/- mice

We previously demonstrated that in a genetic murine models of epilepsy, the Synapsin1-/- mice, eEF2 phosphorylation in brain is strongly increased suggesting that in these mice the pathway that control eEF2 phosphorylation is altered and this might contribute to the excitation/inhibition unbalance and epileptic phenotype (Heise et al., 2017). Thus, we wondered if the level of phosphorylated eEF2 (P-eEF2) is also altered in Scn1a+/- mice.

We analyzed the amount of P-eEF2 phosphorylation in total homogenate of two main brain areas: hippocampus and cerebral cortex. We collected brain samples at different ages (3, 6 and 9 months) in order to detect eventual changes of P-eEF2 in ageing. Western blots analysis showed that the level of P-eEF2 is significantly increased in Scn1a+/- mice at all aged analyzed (with no significant changes among ages) both in One-way ANOVA, Tukey’s post hoc. cortex and in hippocampus (Figure 1A and B) while the level of P-eEF2 was not altered in other tissue such as liver, kidney and heart (Supplementary Figure 1A-C). These data suggest that the specific alteration of the pathway that control eEF2 in brain areas might further contribute to the imbalance of excitation over inhibition occurred in in these mice as we observed in the Synapsin1-/- mice (Heise et al., 2017).

**Figure 1.**
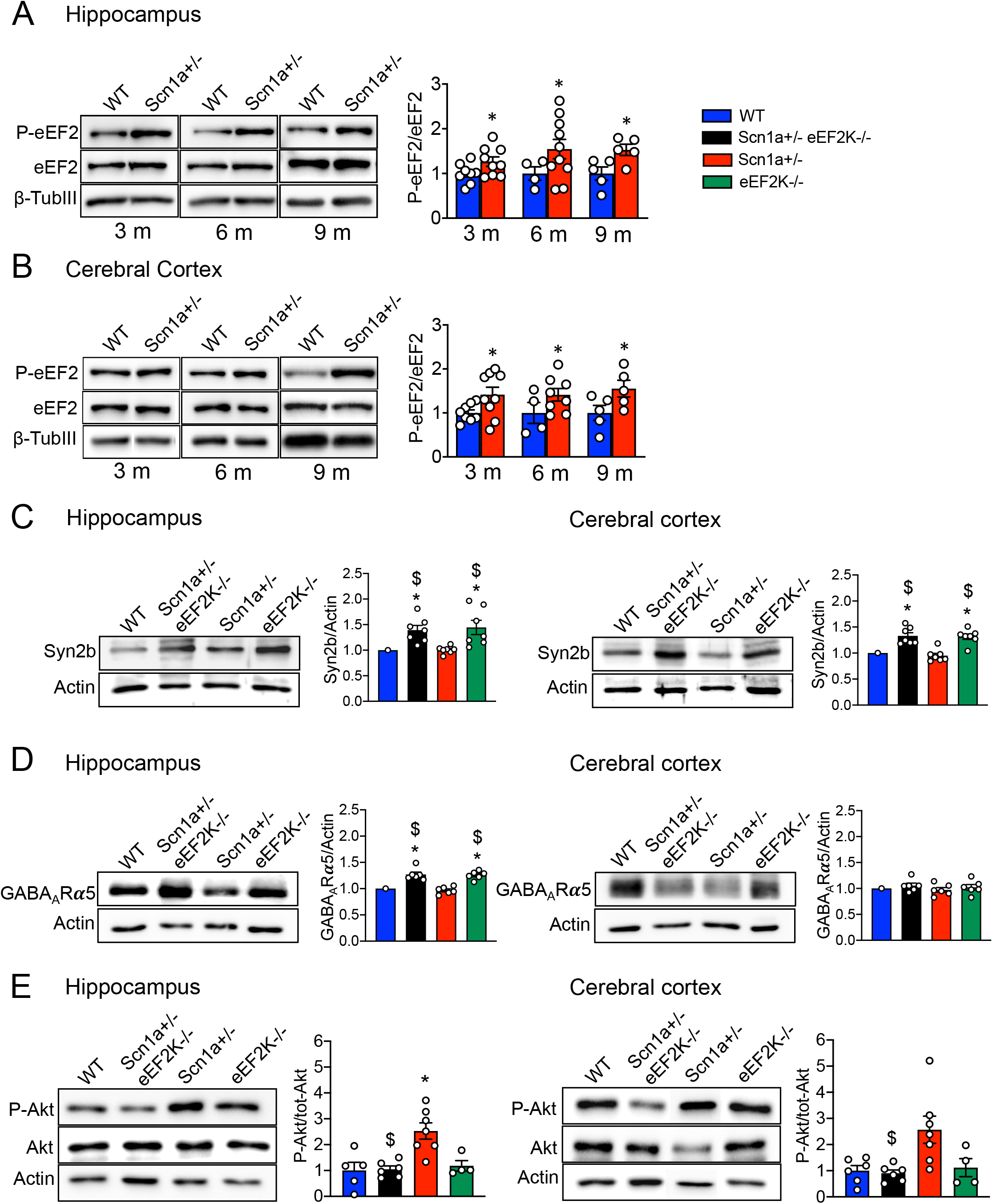
Scn1a+/- mice exhibit higher levels of eEF2 phosphorylation in total homogenate of hippocampus and cerebral cortex compared with WT and eEF2K deletion modulate the expression of a set of proteins that regulate GABAergic synaptic transmission and the level of Akt phosphorylation. **(A-B)** Representative western blots and relative quantification for phosphorylated eEF2 in samples from hippocampus **(A)** and cerebral cortex **(B)** in 3, 6 and 9 months old Scn1a+/- and WT mice. 3 months: hippocampus and cerebral cortex WT n=8, Scn1a+/- n=9; 6 months: hippocampus WT n=4, Scn1a+/- n=10, cerebral cortex WT n=4, Scn1a+/- n=8; 9 months hippocampus and cerebral cortex WT n=5, Scn1a+/- n=5. All Data are presented as mean ± SEM. Statistical analysis *p<0.05 versus corresponding WT; One-sample *t*-test. **(C-D)** Representative western blots and relative quantification show expression levels of synapsin 2b **(C)** and GABA_A_R□5 **(D)** in total homogenate of hippocampus (left) and cerebral cortex (right) from 3 months old WT, Scn1a+/-eEF2K-/-, Scn1a+/- and eEF2K-/- mice. All data are presented as mean ± SEM. N=7 per group for synapsin 2b. N=6 per groups for GABA_A_R□5. Statistical analysis *p<0.05 versus corresponding WT, $p<0.05 versus corresponding Scn1a+/-; One-simple *t*-test. **(E)** Western blots analyses and relative quantification of phosphorylated Akt levels in hippocampus (left) and cerebral cortex (right) of 3 months old WT, Scn1a+/-eEF2K-/-, Scn1a+/- and eEF2K-/- mice. All data are presented as mean ± SEM. WT n=5 (hippocampus) n=6 (cerebral cortex), Scn1a+/- eEF2K-/- n=6, Scn1a+/- n=7, eEF2K-/- n=4. Statistical analysis *p<0.05 versus corresponding WT, $p<0.05 versus corresponding Scn1a+/-; Kruskal-Wallis test, Dunn’s post hoc for hippocampus and One-way ANOVA, Tukey’s post hoc for Cerebral cortex.

### Generation of Scn1a+/-eEF2K-/- mice

To elucidate the role of eEF2K pathway in the etiopathology of Dravet syndrome we deleted the eEF2K gene in Scn1a+/- mice that was previously generated and described in Yu et al. (2006) (Yu et al., 2006) and mimic the Dravet syndrome described in human patients. We crossed Scn1a+/- mice with eEF2K-/- mice (generated by the laboratory of Prof. Alexey G. Ryazanov, Rutgers University) and as expected by the Mendelian law, in the first generation we obtained about 50% of the mice with the genotype Scn1a+/+ eEF2K+/- and 50% with the genotype Scn1a+/- eEF2K+/-.

We then crossed the Scn1a+/- eEF2K+/- mice in order to obtained the three main genotypes needed for our studies: Scn1a+/+eEF2K+/+ (WT mice), Scn1a+/-eEF2K+/+ mice (Scn1a+/- mice) and Scn1a+/-eEF2K-/- mice (Supplementary Figure 2A). We also obtained Scn1a+/+eEF2K-/- mice (eEF2K KO mice) that were used to study if eEF2K deletion caused any behavioral alterations. To avoid genetic background influence, well described in the Scn1a+/- mice (Rubinstein et al., 2015; Kang et al., 2019), the mice were backcrossed for about 20 generations before were used for all the experiments in order to obtain the same genetic background among genotypes. As expected the level of P-eEF2 was completely abolished in the Scn1a+/-eEF2K-/- mice (Supplementary Figure 2B).

### eEF2K deletion modulate the expression of a set of proteins that regulate GABAergic synaptic transmission and the level of Akt phosphorylation

We previously showed that in eEF2K-/- mice the increased efficiency of GABAergic transmission was mediated by higher levels of expression of Synapsin 2b (Syn2b) in cortex and hippocampus and α5 subunit containing GABA_A_ receptor (GABA_A_R□5) in hippocampus (Heise et al., 2017). We thus analyzed the expression of Syn2b and GABA_A_R□5 subunit in Scn1a+/- and Scn1a+/-eEF2K-/- mice and we confirmed a significant increased expression of both Syn2b and α5-GABAA in hippocampus and of Syn2b in cortex in mice deleted of eEF2K (eEF2K-/- and Scn1a+/-eEF2K-/- mice). On the contrary the expression of both Syn2b and GABA_A_R□5 was not changed in the Scn1a+/- mice (Figure 1C and D). All these data suggest that deletion of eEF2K was able to strength inhibitory synapses in Scn1a+/- mice.

Moreover, alteration of the Akt signaling has been also recently described in the Scn1a+/- mice (Tai et al., 2020). Interestingly our results showed that the increased levels of P-Akt in hippocampus and cortex of Scn1a+/- mice were rescue to level similar to WT mice in the Scn1a+/- mice by the deletion of eEF2K (Figure 1E).

### eEF2K deletion protects Dravet mice from epileptic seizures onset

We first investigate the outcome of eEF2K deficiency on epileptic symptoms displayed by the Scn1a+/- mice. We recorded electroencephalographic (EEG) traces from male mice for 24 hours and we counted the number of spikes that Scn1a+/- mice and Scn1a+/-eEF2K-/- mice displayed compared to WT mice. While Scn1a+/- mice displayed an increased number of spikes in basal condition compared to WT mice, the Scn1a+/-eEF2K-/- mice showed an EEG traces not different from WT mice, suggesting that eEF2K depletion was able to rescue the altered EEG observed in the Scn1a+/- mice (Figure 2A left panels and B). In Dravet syndrome, the onset of the epileptic seizures is triggered by an episode of fever and, usually, the following seizures arise without thermal stimuli (Jansen et al., 2006). This feature is maintained in Scn1a+/- mice (Oakley et al., 2009), therefore we triggered the convulsions by increasing mice body temperature and we counted the number of spikes 7 days after. We found a strong increase of number of spikes measured in 24 hours in Scn1a+/- mice after the thermal stress but not in the Scn1a+/-eEF2K-/- mice, suggesting that eEF2K deficiency reduced the susceptibility to epilepsy of the Scn1a+/- mice (Figure Fig. 2A right panels and B). Furthermore, eEF2K depletion significantly increased the minimum temperature necessary to trigger epileptic seizures (Figure 2C) and the time before the first seizure appear upon the body temperature reach 40°C in comparison to Scn1a+/- mice (Figure 2D)

**Figure 2.**
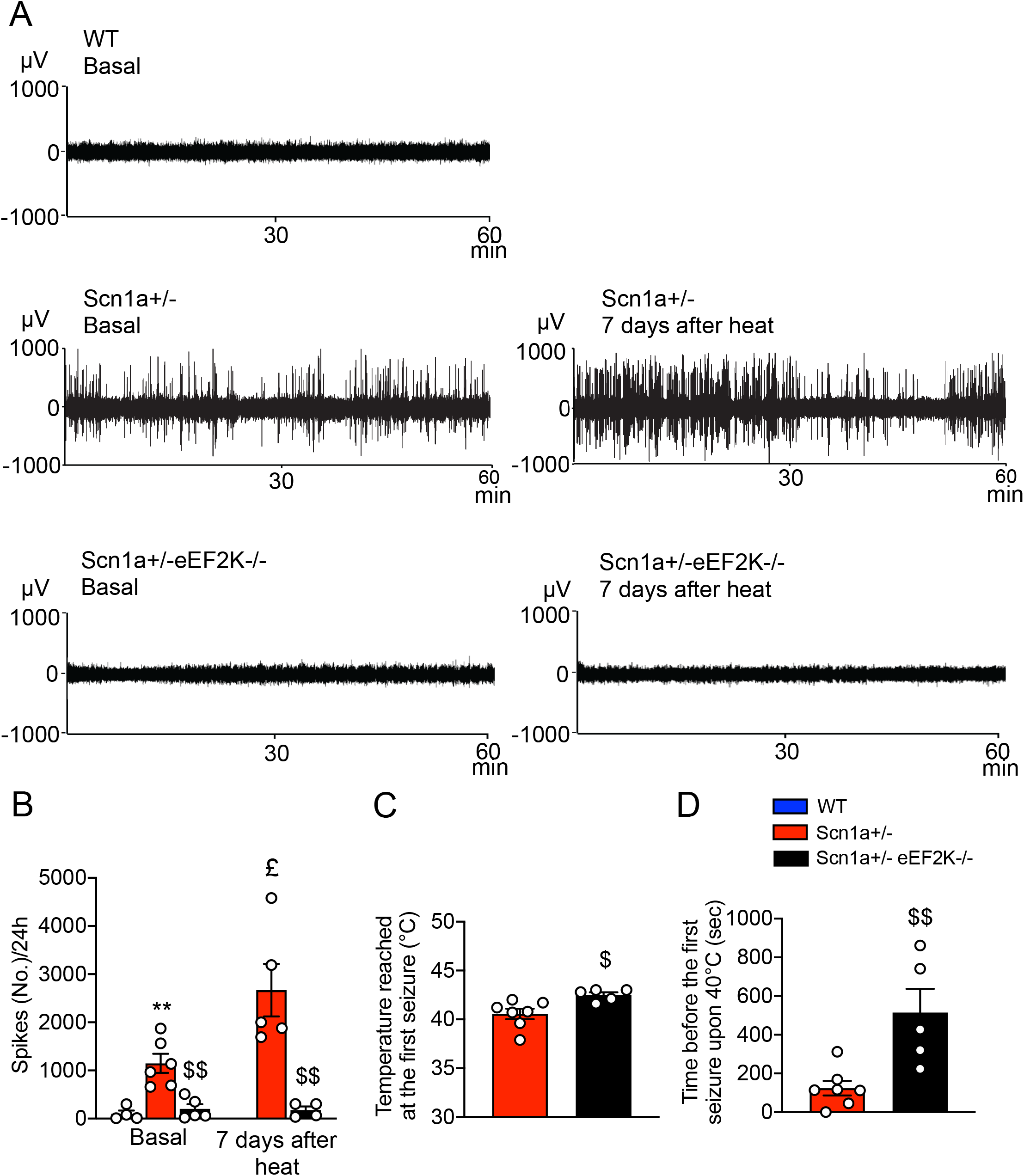
eEF2K deletion protect Scn1a+/- mice from epileptic seizure onset. **(A)** Representative EEG traces (a representative 60 min. registration is shown) of a WT, Scn1a+/- and Scn1a+/-eEF2K-/- mice in basal condition (left) and 7 days after thermal stress (right). Scn1a+/- mice show high number of EEG spikes in basal and 7 days after thermal stress condition when compared with WT and Scn1a+/-eEF2K-/- mice. **(B)** Quantification of the number of EEG spikes per 24h in basal condition (WT n=4, Scn1a+/- n=6 and Scn1a+/- eEF2K-/- n=5) and 7 days after heat (Scn1a+/- n=5 and Scn1a+/- eEF2K-/- n=4). Scn1a+/-eEF2K-/- mice are clearly less susceptible to seizure in basal and 7 days after thermal stress condition than Scn1a+/- mice. Data are presented as mean ± SEM. Statistical analysis for number of EEG spikes in basal condition **p<0.01 versus corresponding WT, $$p<0.01 versus corresponding Scn1a+/-; One-way ANOVA test, Tukey’s post hoc. Statical analysis for number of EEG spikes 7 days after thermal stress condition $$p<0.01 versus corresponding Scn1a+/-; Unpaired two-tailed *t*-test. Statistical analysis £p<0.05 versus corresponding Scn1a+/- in basal condition; Unpaired two-tailed *t*-test. **(C)** Temperature reached at the first seizure. eEF2K deletion increase the minimum temperature necessary to reach the first seizure in Scn1a+/- mice. Data are presented as mean ± SEM. Scn1a+/- n=7, Scn1a+/- eEF2K-/- n=5. Statistical analysis $p<0.05 versus corresponding Scn1a+/-; Unpaired two-tailed *t*-test. **(D)** Time before the first seizure occur at 40°C. eEF2K deletion increase the time before the first seizure occurred or until 42.5°C was reached. Data are presented as mean ± SEM. Scn1a+/- n=7, Scn1a+/- eEF2K-/- n=5. Statistical analysis $$p<0.01 versus corresponding Scn1a+/-; Unpaired two-tailed *t*-test.

### eEF2K deletion enhances GABAergic transmission in Scn1a+/- mice

A hyperexcitable neuronal network is one of the major causes of the epileptic seizures in Dravet syndrome (Liautard et al., 2013; Hedrich et al., 2014). Given that in the eEF2K-/- mice GABAergic transmission is potentiated (Heise et al., 2017), we hypothesized that the Scn1a+/-eEF2K-/- mice were less susceptible to develop epileptic seizure because of the potentiation of their GABAergic synaptic transmission. To verify this hypothesis, we recorded spontaneous inhibitory postsynaptic currents (sIPSC) in pyramidal neurons of the CA1 region of the hippocampus in brain slices from WT, Scn1a+/- and Scn1a+/-eEF2K-/-mice. As expected, Scn1a+/- mice displayed reduced amplitude and instantaneous frequency in the recorded sIPSCs compared to WT mice (Yu et al., 2006). Interestingly eEF2K depletion enhanced both amplitude and instantaneous frequency of sIPSC of Scn1a+/- mice. However, the rescue was partial, indeed the iPSC amplitude and instantaneous frequency measured in Scn1a+/-eEF2K-/- were significantly smaller compare to WT mice (Figure 3A-C).

**Figure 3.**
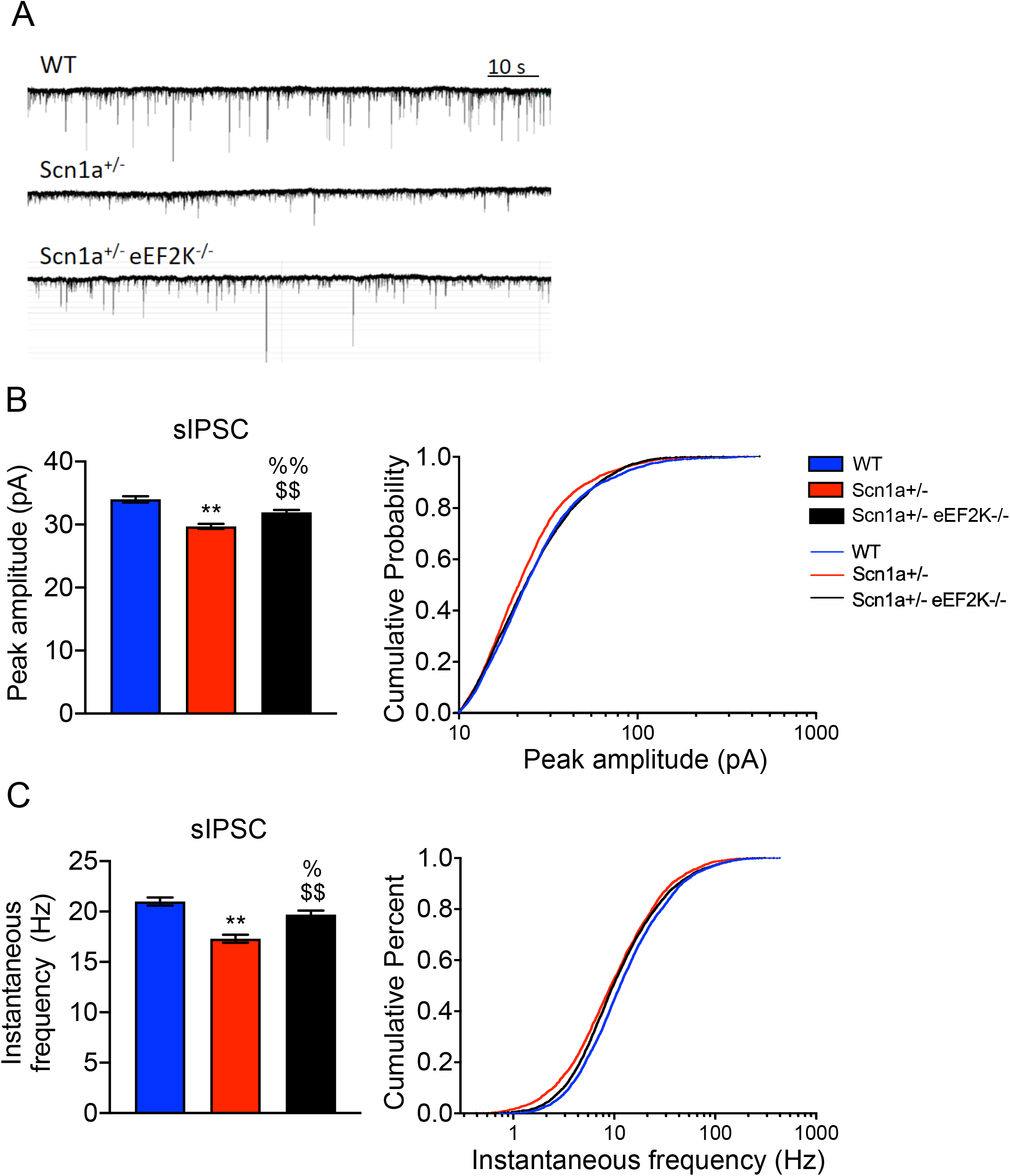
eEF2K deletion enhances GABAergic transmission in Scn1a+/- mice. **(A)** Representative sIPSC trace measured in hippocampal neurons of CA1 region of WT, Scn1a+/- and Scn1a+/-eEF2K-/- mice. **(B-C)** Quantification of peak amplitude **(B)** and instantaneous frequency **(C)** from sIPSC traces. All data are presented as mean ± SEM. Statistical analysis for peak amplitude **p<0.001, %% p<0.05 versus corresponding WT; $$ p<0.001 versus Scn1a+/-; One-way ANOVA, Bonferroni. Statistical analysis for instantaneous frequency **p<0.001, %p<0.01 versus corresponding WT; $$ p<0.05 versus Scn1a+/-; One-way ANOVA, Bonferroni.

### eEF2K deletion ameliorates motor coordination in Scn1a+/- mice

Motor deficits are common in Dravet syndrome: most of the patient are affected by ataxia, dysarthria, intention tremor and eye movement disorder (Genton et al., 2011) and these features are replicated in Scn1a+/- mice (Yu et al., 2006; Han et al., 2012). We thus tested WT, Scn1a+/- and Scn1a+/-eEF2K-/- mice for motor function with the spontaneous motor activity test, the limb force with the wire hanging test and the motor coordination with balance beam, pole test and rotarod test. As previously demonstrated by Heise et al. (2017), eEF2K-/- do not show alteration in motor function and limb force when compared with WT mice. First, we verified the effect of eEF2K deletion on motor coordination comparing eEF2K-/- mice with WT mice. We recorded horizontal and vertical movements of freely moving animals for 3 hours. As showed in the supplementary Figure 3 motor coordination were also not altered in the eEF2K-/- mice (Supplenetary Figure 3A-C).

WT and Scn1a+/- mice were then compared to the Scn1a+/-eEF2K-/- mice. Movements in horizontal and vertical directions were not statistically different, meaning that Scn1a+/- mice had no difficult in walking or climbing. Also Scn1a+/-eEF2K-/- mice didn’t display difficulty in movements, even if vertical movements were slightly significantly higher compared to Scn1a+/- mice (Figure 4A). We also measure the latency to fall in wire hanging test as measure of limb muscle strength. We did not fin any difference in terms of muscle strength among the three groups of animals (Figure 3B).

**Figure 4.**
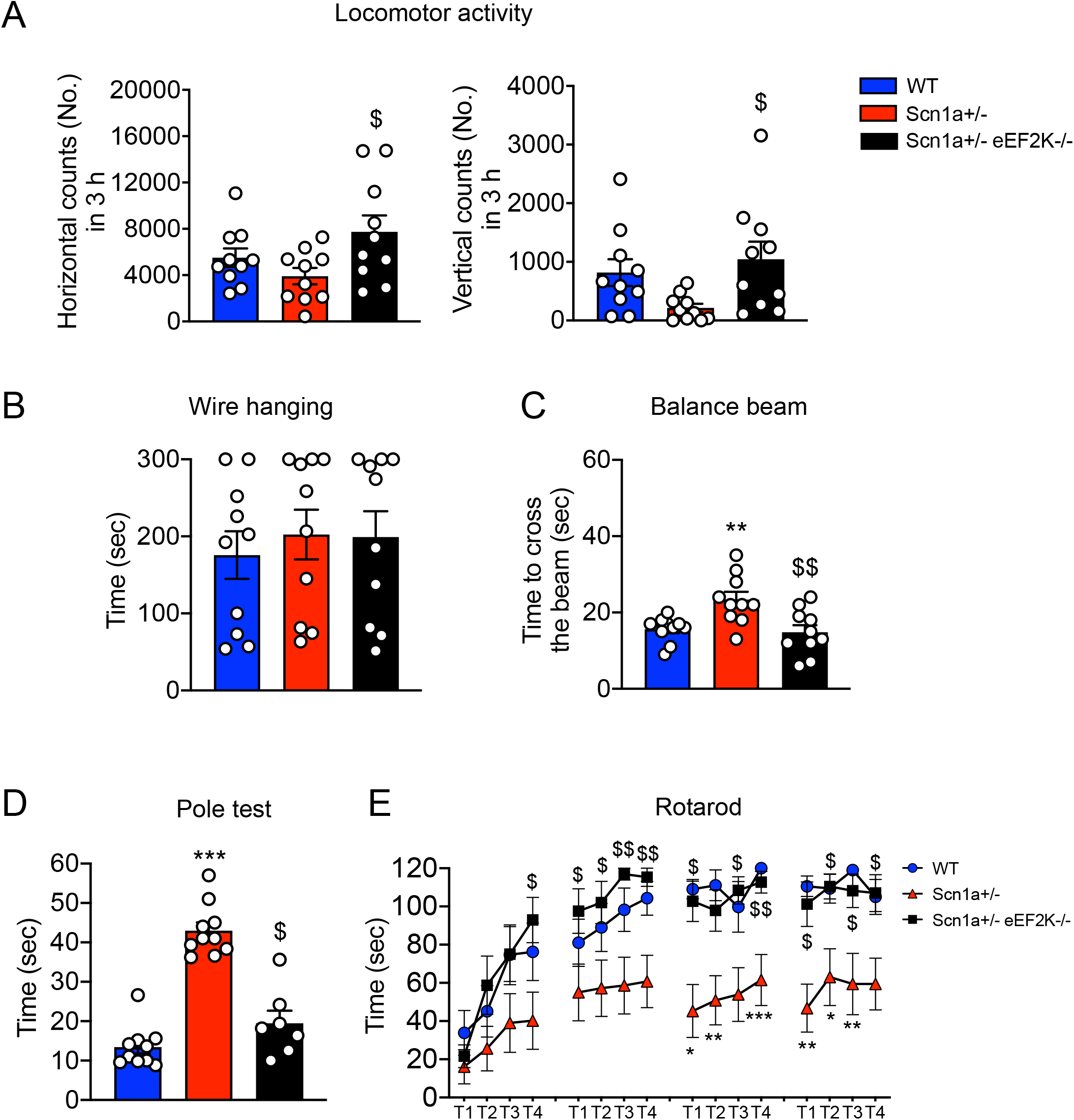
eEF2K deletion rescues balance impairment and motor coordination in Scn1a+/- mice. **(A)** Counting of horizontal (left) and vertical (right) movements occurred in 3 hours in the locomotory activity test. Scn1a+/-eEF2K-/- mice show high number of horizontal and vertical movements. Scn1a+/- had no difficulty in movement or climbing. Data are presented as mean ± SEM. N=10 per groups. Statistical analysis $p<0.05 versus corresponding Scn1a+/-; One-way ANOVA, Tukey’s post hoc. **(B)** Time to fall in wire hanging test measured over 300 seconds. Data are presented as mean ± SEM. N=10 per groups. One way-ANOVA, Tukey’s post hoc. **(C)** Time to cross the 12 mm beam of trained WT, Scn1a+/- and Scn1a+/-eEF2K-/- mice. Scn1a+/- mice are impaired in the balance beam test when compared with WT and Scn1a+/-eEF2K-/- mice. All data are presented as mean ± SEM. N=10 per groups. Statistical analysis **p<0.01 versus corresponding WT, $$p<0.01 versus corresponding Scn1a+/-; One way-ANOVA, Tukey’s post hoc. **(D)** Time to downstream the pole in pole test. Scn1a+/- are impaired in the pole test when compared with WT and Scn1a+/-eEF2K-/- mice. All data are presented as mean ± SEM. WT n=10, Scn1a+/- n=10, Scn1a+/- eEF2K-/- n=7. Statistical analysis ***p<0.001 versus corresponding WT; $p<0.05 versus Scn1a+/-; Kruskal-Wallis test, Dunn’s post hoc. **(E)** Scn1a+/- mice show impaired motor coordination in rotarod test when compared with WT and Scn1a+/-eEF2K-/-. The plot shows mean ± SEM latency to fall from an accelerating rotarod during trials. N=10 per group. Statistical analysis *p<0.05, **p<0.01, ***p<0.001 versus corresponding WT; $p<0.05, $$p<0.01 versus Scn1a+/-; Kruskal-Wallis test, Dunn’s post hoc.

We then assayed the motor coordination with balance beam and we found that while the Scn1a+/- mice took longer time compare WT mice to cross the 12 mm bean the Scn1a+/-eEF2K-/- mice behaved as the WT mice (Figure 4C). With the pole test we measured the latency to turn downwards and descend and we observed that the Scn1a+/- mice took longer time than WT mice while the Scn1a+/-/eEF2K-/- mice behaved as WT mice (Figure 4D). Finally, in the rotarod test the Scn1a+/- mice showed a strong impairment in motor coordination that was rescued in the Scn1a+/- eEF2K-/- mice (Figure 4E). All these data clearly demonstrated that deletion of eEF2K was able to recover all the motor deficits found in the Scn1a+/- mice.

### eEF2K deletion rescue the stereotyped behavior of Scn1a+/- mice

Dravet syndrome patients are affected also by important and diverse patient-related co-morbidity such as stereotyped behaviors and anxiety

We measured stereotyped behaviors by self-grooming test, counting the time spent in this activity. We observed that EF2K deletion in Scn1a+/- mice was able to reduce the time occupied in self-grooming of Scn1a+/- mice to value similar to the WT mice (Figure 5A). As expected the eEF2K-/- mice did not show any increased grooming activity compare to WT mice (Supplementary Figure 4A).

**Figure 5.**
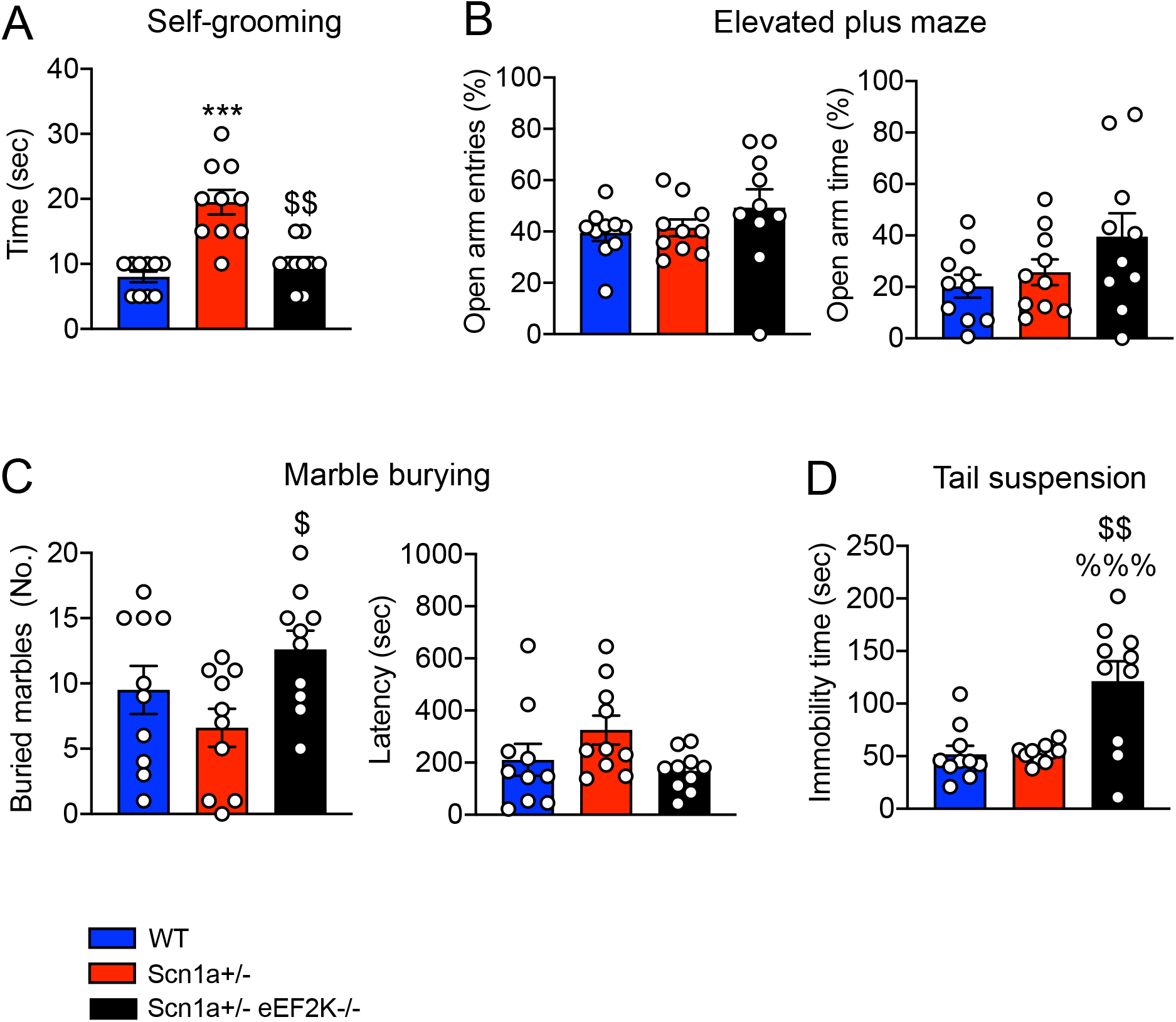
Genetic deletion of eEF2K rescue autistic-like behaviour in Scn1a+/- mice. **(A)** Significantly increased in repetitive grooming behavior in Scn1a+/- mice compared with WT and Scn1a+/-eEF2K-/- evaluated in terms of time spent to do grooming over 10 min observation. Data are presented as mean ± SEM. N=10 per group. Statistical analysis ***p<0.001 versus corresponding WT; $$p<0.01 versus corresponding Scn1a+/-; Kruskal-Wallis test, Dunn’s post hoc. **(B-C)** Scn1a+/- mice do not show anxiety-behavior in elevated plus maze **(B)** and marble burying **(C)** test. **(B)** Open arm entries (left) and time spend in open arms (right), evaluated in the elevated plus maze test over 5 minutes. Data are presented as mean ± SEM. N=10 per group. One-way ANOVA, Tukey’s post hoc. **(C)** Number of marbles buried (left) and latency to the first buried (right) evaluated in the marble-burying test over 15 min. Data are presented as mean ± SEM. N=10 per group. Statistical analysis $p<0.05 versus corresponding Scn1a+/-; One-way ANOVA, Tukey’s post hoc. **(D)** Time spent immobile in tail suspension test do not show depressive-like behaviours in Scn1a+/- mice when compared with WT. Scn1a+/-eEF2K-/- mice show depressive-like behaviour when compared with WT and Scn1a+/- mice. Data are presented as mean ± SEM. N=10 per group. Statistical analysis %%%p<0.001 versus corresponding WT, $$p<0.01 versus corresponding Scn1a+/-; One-way ANOVA, Tukey’s post hoc.

Then we tested mice for anxiety-like behavior. We did not found difference among the three genotypes nor in the elevated plus maze nor in the marble burying test demonstrating that eEF2K deletion did not affect anxiety-like behavior (Figure 5B and C), as also previously demonstrated by Heise et al. (2017) where eEF2K-/- mice behave like WT when tested for anxiety-like behavior. On the contrary when we analyzed depression like behavior by using the tail suspension test we found that eEF2K deletion was associated to an increased immobility time in the tail suspension test, both when deleted in the WT mice (supplementary Figure 4B) and in the Scn1a+/- mice (Figure 5D).

### eEF2K deficiency ameliorates episodic and spatial memory and social novelty preference of Scn1a+/- mice

Another typical trait of Dravet patients is the presence of cognitive impairments (Jansen et al., 2006; Dravet, 2011b). We tested mice for episodic and spatial memory using novel object recognition test and spatial object recognition test, respectively. Mice had to recognize the unfamiliar or displaced object after 5 minutes, 120 minutes or 24 hours from familiarization phase. In the novel object recognition test Scn1a+/- mice were unable to recognize the new object as unfamiliar, as shown by the reduced discrimination index at all the time points compared to WT mice, suggesting that Scn1a+/- mice had deficits in episodic memory. Interestingly the Scn1a+/- eEF2K-/- mice, as well as WT mice, displayed a strong preference for the unfamiliar objects, implicating that eEF2K deletion had a beneficial effect on Scn1a+/- mice episodic memory (Figure 6A). Compared to WT mice the eEF2K-/- mice did not showed any impairment (Supplementary Figure 4C and D).

**Figure 6.**
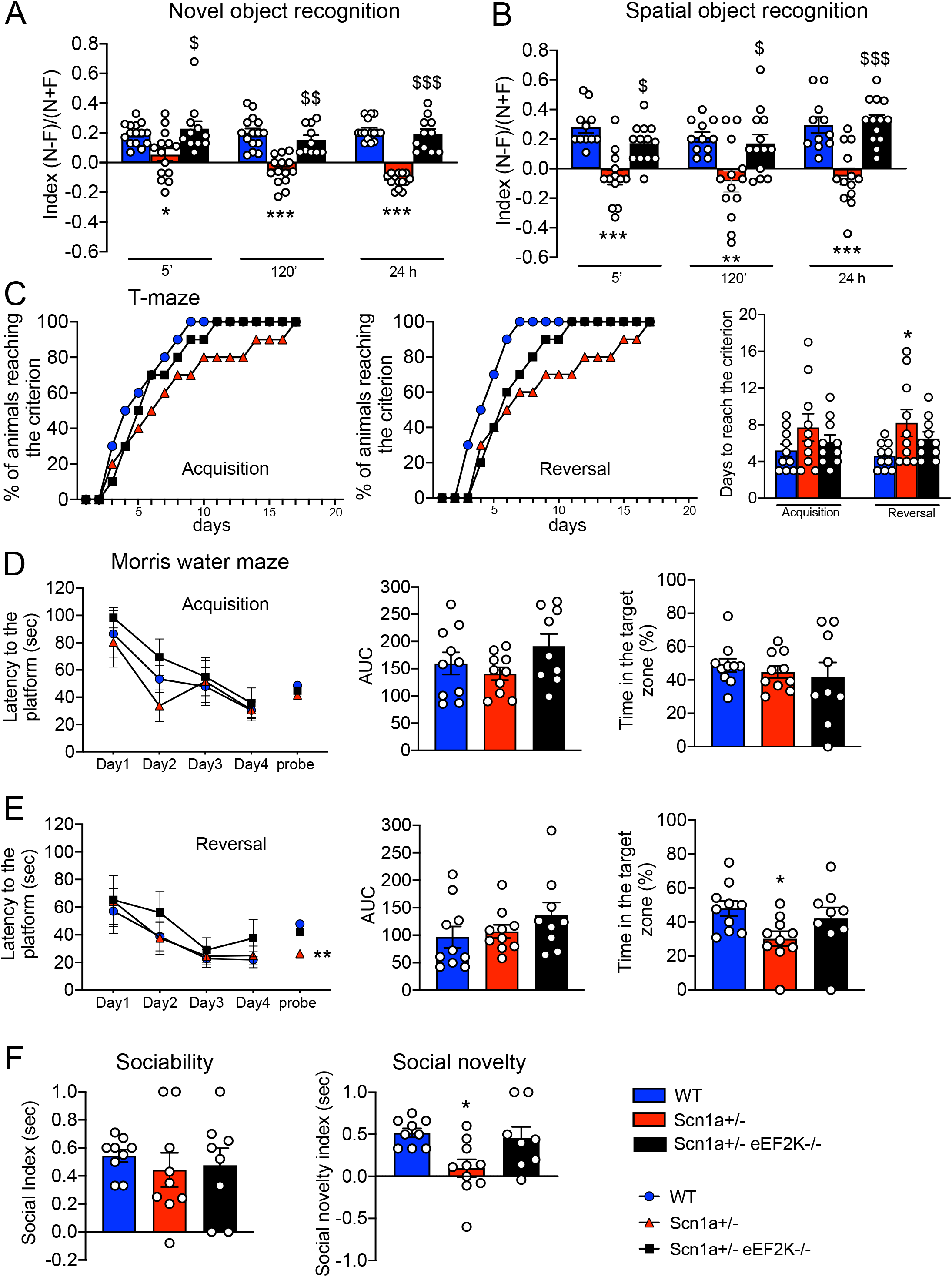
eEF2K deletion in Scn1a+/- mice rescues the learning and memory impairment and social recognition alteration. **(A-B)** Discrimination index evaluated in the novel object recognition test **(A)** and in the spatial object recognition test **(B). (A)** Scn1a+/- mice show an impairment in novel object recognition at 5 min, 120 min and 24h after the familiarization phase compared with WT and Scn1a+/-eEF2K-/- mice. Data are presented as mean ± SEM. 5 min and 24 h: WT n=15, Scn1a+/- n=14, Scn1a+/-eEF2K-/- n=11; 120 min WT n=15 Scn1a+/- n=14, Scn1a+/-eEF2K-/-=10. Statistical analysis *p<0.05, ***p<0.001 versus corresponding WT, $p<0.05, $$p<0.01, $$$p<0.001 versus corresponding Scn1a+/-; Kruskal-Wallis test, Dunn’s post hoc. **(B)** Scn1a+/- mice are impaired in spatial object recognition tested at 5 min, 120 min and 24h after familiarization phase when compared with WT and Scn1a+/-eEF2K-/- mice. Data are presented as mean ± SEM. WT n=11, Scn1a+/- n=13, Scn1a+/-eEF2K-/- n=13. Statistical analysis **p<0.01, ***p<0.001 versus corresponding WT, $p<0.05, $$$p<0.001 versus corresponding Scn1a+/-; Kruskal-Wallis test, Dunn’s post hoc for 5 min and One-way ANOVA, Tukey’s post hoc for 120 min and 24h. **(C)** Scn1a+/- mice required more days to achieve the criterion than WT mice during the reversal phase of T-maze. Data are presented as mean ± SEM. N=10 per group. Statistical analysis *p<0.05 versus corresponding WT. Days to reach the criterion was analyzed by One-way ANOVA, Tukey’s post hoc. The percentage of animals reaching the criterion was analyzed by Fisher’s exact test. **(D-E)** Acquisition and reversal learning in Morris Water Maze task in terms of latency to reach the platform across 4 days trial and probe test and in percentage of the time spent in the target quadrant during the probe test. **(D)** Scn1a+/- mice display normal Morris water maze learning in the acquisition phase in both latency to reach the platform and in probe test. Data are presented as mean ± SEM. WT, Scn1a+/- n=10, Scn1a+/-eEF2K-/- n=9. Each day latency was analyzed by Kruskal-Wallis test, Dunn’s post hoc. AUC and percentage in the target zone was analyzed by One-way ANOVA, Tukey’s post hoc. **(E)** Scn1a+/- mice show an impaired spatial memory in Morris water maze probe test in the reversal phase. Data are presented as mean ± SEM. WT, Scn1a+/- n=10, Scn1a+/-eEF2K-/- n=9. Each day of latency was analyzed by Kruskal-Wallis test, Dunn’s post hoc. Statistical analysis in reversal probe test latency **p<0.01 versus corresponding WT; One-way ANOVA, Tukey’s post hoc. AUC was analyzed by Kruskal-Wallis test, Dunn’s post hoc. Statistical analysis for percentage of time in the target zone *p<0.05 versus corresponding WT; One-way ANOVA, Tukey’s post hoc. **(F)** Sociability index (left) and social novelty preference index (right) evaluated in sociability (left) and in social novelty test (right). Scn1a+/- mice display a normal social attitude when compared with WT and Scn1+/-eEF2K-/- mice in sociability test (left). Data are presented as mean ± SEM. WT, Scn1a+/- n=9, Scn1a+/-eEF2K-/- n=8; One-way ANOVA, Tukey’s post hoc. Scn1a+/- display an impairment in social recognition when tested for social novelty test (right). Data are presented as mean ± SEM. WT n=9, Scn1a+/- n=10, Scn1a+/-eEF2K-/- n=8; One-way ANOVA, Tukey’s post hoc.

In the spatial object recognition test Scn1a+/- mice displayed a defect preferring the stationary object, as shown by the negative discrimination index. On the contrary Scn1a+/-eEF2K-/- performed in the test similarly to the WT mice spending more time exploring the displaced object (Figure 6B) confirming the positive effect of eEF2K deletion on memory performances. In the T-maze and Morris water maze tests, two other memory related test, the Scn1a+/- mice performed as the WT mice in the acquisition phase (Figure 6C left and right and D) while showed a significant impairment in the reversal phase of the test (Figure 6C center and right, E) that was rescued by the deletion of eEF2K (Figure 6C, E).

We then analyzed the social behavior of WT, Scn1a+/- and Scn1a+/-eEF2K-/- mice using the sociability and social novelty preference test. Scn1a+/- mice exhibited more interest for the unfamiliar mouse than for the empty cage, as well as WT and Scn1a+/-eEF2K-/- mice, demonstrating a normal attitude in social interaction. However, in the social novelty test, Scn1a+/- mice, differentially to WT mice, spent the same time in exploring familiar and unfamiliar mice, indicating impairment in the social recognition. Interestingly the deletion of eEF2K in the Scn1a+/- was able to rescue this behavioral alteration (Figure 6F). Indeed the eEF2K-/- mice were not different from WT mice for both social novelty test (Supplementary Figure 4G) and sociability (Heise et al., 2017).

Finally we also observed that Scn1a+/- mice, were impaired in both olfactory and visual cliff tests and that eEF2K deletion was able to rescue these behavioral alterations (Supplementary Figure 5A and B).

### Pharmacological rescue of EEG alterations of Scn1a+/- mice using an eEF2K inhibitor

We demonstrated that the genetic deletion of eEF2K in the Scn1a+/- mice rescued the main phenotypes such as epileptic seizure and deficit in behaviour, which characterized the Dravet syndrome suggesting eEF2K as a possible pharmacological target.

To confirm our hypothesis, we tested the ability of A484954 of reverting EEG alterations found in Scn1a+/- mice. A484954 is a selective small molecule inhibitor for eEF2K that inhibits eEF2K activity in an ATP-competitive but not CaM-dependent manner (Kodama et al., 2019), resulting in an inhibition of eEF2 phosphorylation at Thr 56. To delivery A484954 in mice we used the pellet implantation methodology provided by Innovative Research of America company (IRA), instead of classical intraperitoneal injection.

Mice were implanted with the pellets on the lateral side of the neck between the ear and the shoulder. We used 10mg of A484954/pellets, which released 1 mg of drug every day for 10 days. Placebo pellets were used in the experiment as the proper control. The treatment was performed as described in Figure 7A. First, we confirmed a significant reduced phosphorylation of eEF2 in the hippocampus and cerebral cortex of Scn1a+/- treated with A484954 mice in comparison with the ones treated with placebo after 10 days of treatment, confirming that A484954 was able to inhibit eEF2K activity in brain (Figure 7B).

**Figure 7.**
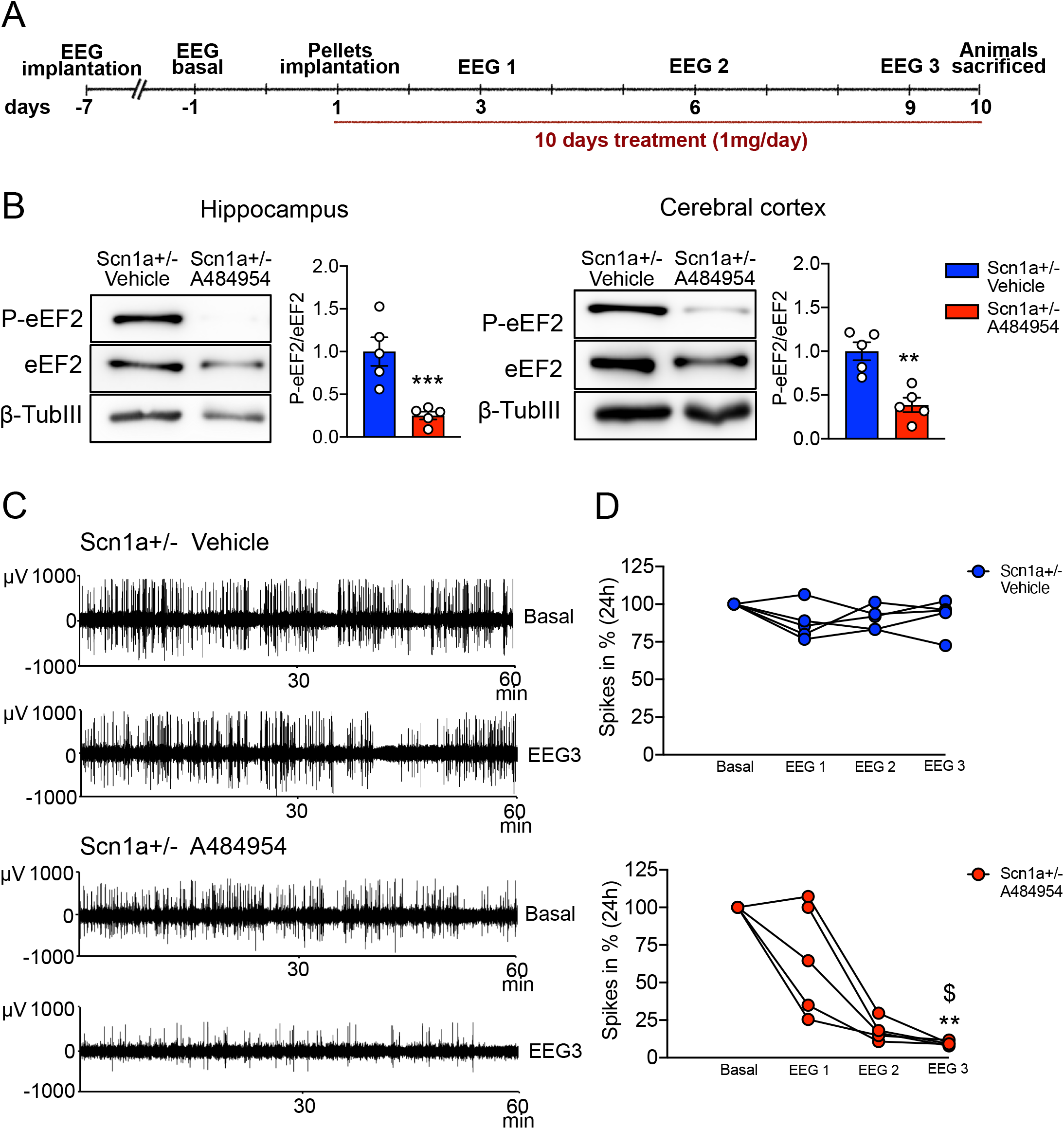
Pharmacological inhibition of eEF2K activity rescue the epileptic phenotype in Scn1a+/- mice. **(A)** Experimental timeline. **(B)** Representative western blots and relative quantification show a significantly reduction of phosphorylated eEF2 in total homogenate of hippocampus (left) and cerebral cortex (right) of Scn1a+/- mice after a chronic treatment of 10 days with 1mg/day dose of A484954 compared with vehicle. All data are presented as mean ± SEM. N=5 per group. Statistical analysis ***p<0.001, **p<0.01 versus corresponding Scn1a+/- treated with vehicle; One-simple *t*-test. **(C)** Representative EEG traces (60 min registration is shown) in basal condition and at 9 days of treatment (EEG3) of a Scn1a+/- mice treated with vehicle or A484954. **(D)** Quantification of the number of EEG spikes calculated in percentage per 24h in basal condition and at 3 (EEG1), 6 (EEG2) and 9 (EEG3) days of treatment. Significant reduction of the number of EEG spikes across the three EEG recording in Scn1a+/- mice treated with A484954. No difference in number of EEG spikes in Scn1a+/- mice treated with vehicle. Scn1a+/- treated with A484954 n=5, Scn1a+/- treated with vehicle n=5. Statistical analysis **p<0.01 versus corresponding basal, $p<0.05 versus corresponding EEG1; Friedman test.

We then measure the EEG number of spikes in the same mice before pellet implantation (EEG basal), and after 3, 6 and 9 days (EEG1, EEG2 and EEG3) of treatment. We observed a progressive decrease of number of spikes in the Scn1a+/- mice treated with A484954 (Figure 7C and D) demonstrating that also pharmacological inhibition of eEF2K was able to strongly reduce the number of EEG spikes, and probably to ameliorate the epileptic phenotype, of the Scn1a+/- mice.

## Discussion

We have previously identified eEF2K, a kinase involved in the regulation of protein translation, as an effective modulator of GABAergic synaptic transmission (Heise et al., 2017). Indeed, we demonstrated that genetic and pharmacological eEF2K inhibition rescued the epileptic phenotype in a genetic animal model of epilepsy, the Synapsin1 knock out mice (Heise et al., 2017). With this work we provide strong evidence that genetic deletion of eEF2K in the Scn1a+/- mice, a mouse model of Dravet syndrome, can rescue both epileptic phenotype and behavioral alterations.

The function of eEF2K in regulating synaptic plasticity and the excitation/inhibition balance in the brain has been widely studied using in vitro and in vivo models, suggesting that eEF2K dysregulation may contribute to the pathogenesis of a subset of neurodegenerative and neuropsychiatric disorders (Monteggia et al., 2013; Jan et al., 2018; Beckelman et al., 2019; Beretta et al., 2020) and making the eEF2K/eEF2 pathway a valuable pharmacological target for brain related disease treatment. High levels of phosphorylated eEF2 have been described in Alzheimer disease (Ma et al., 2014; Jan et al., 2017), Parkinson disease (Jan et al., 2018) and genetic epilepsy (Heise et al., 2017) and it has also been reported that inhibition of eEF2K activity ameliorate the pathology (Ma et al., 2014; Heise et al., 2017; Jan et al., 2017; Beckelman et al., 2019).

We found that phosphorylation of eEF2, the only substrate of eEF2K, is higher in the hippocampus and cerebral cortex of Scn1a+/- mice compared with WT, while no difference was found in the liver, kidney and heart. We measured a higher level of phosphorylated eEF2 in adult Scn1a+/ mice (3 months old mice) when the high number of spikes and the behavioral deficit are clearly displayed. The increased level of eEF2 phosphorylation was maintained also in 9 month old mice. These data suggest eEF2K/eEF2 pathway alterations might contribute to the inhibition/excitation unbalance that leads to epileptic phonotype and behavioral deficit of Scn1a+/- mice. Indeed, even if the molecular mechanism responsible for the increased level of eEF2 phosphorylation in the Scn1a+/- mice remain to be determined, we demonstrated that the genetic deletion of eEF2K was sufficient to strongly reduce the interictal activity in basal and under thermal stress conditions. As previously reported in the literature (Yu et al., 2006; Mantegazza et al., 2021), mutations in the Nav1.1 channel caused selective reduction in sodium currents and action potential firing of GABAergic interneurons in hippocampus neocortex, cerebellum and thalamus in Scn1a+/- mice leading to reduced GABAergic transmission (Yu et al., 2006; Kalume et al., 2007; Han et al., 2012; Hedrich et al., 2014; Tai et al., 2014). Given that eEF2K-/- mice exhibited a stronger GABAergic synaptic transmission (Heise et al., 2017), we asked whether this deficit in GABAergic transmission in Scn1a+/- could be reverted by eEF2K deletion. We found that eEF2K deletion in Scn1a+/- mice enhanced both amplitude and instantaneous frequency of sIPSCs in CA1 pyramidal neurons. Moreover, we demonstrated that eEF2K deletion specifically potentiated the synaptic and tonic GABAergic transmission increasing the expression of Synapsyn2b (Medrihan et al., 2013) and GABA_A_R□5 (Scimemi et al., 2005; Glykys et al., 2008) respectively. In conclusion these data suggest that eEF2K deletion is sufficient to rescue the excitation/inhibition unbalance responsible for the epileptic phenotype of Scn1a+/- mice by selectively increasing GABAergic transmission efficiency.

Several data have demonstrated that PI3K/Akt/mTOR signaling plays an important role in synaptic plasticity and may influence neurological excitability (Meng et al., 2013; Mossa et al., 2021; Sharma and Mehan, 2021). It has also been demonstrated that the PI3K/Akt/mTOR pathway is overactivated in different ASD and epilepsy mouse models including the Scn1a+/- mouse model (Xing et al., 2019; Tai et al., 2020). We found that in these mice the increased level of phosphorylated Akt was rescued by the deletion of eEF2K. Given the complexity of PI3K/Akt/mTOR and eEF2K/eEF2 pathway interconnections in the regulation in neuronal plasticity (Wang et al., 2001; Heise et al., 2014), our data suggest the eEF2K deletion might have a therapeutic antiepileptic effect on Scn1a+/- mice by not only directly modulating GABAergic transmission but also acting at the level of PI3K/Akt/mTOR pathway. However, the interplay between these pathways in the context of epilepsy and ASD and how the deletion of eEF2K restored normal level of phosphorylated Akt needs further investigation.

Because it has been demonstrated that the eEF2/eEF2K signaling pathway is implicated in memory formation and synaptic function, and its inhibition rescued memory deficits in aged mice (Gosrani et al., 2020) and improved adult neurogenesis in the dentate gyrus (DG) of the hippocampus (Taha et al., 2020), we investigated the effect of eEF2K deletion on the motor and behavioral deficits of the Scn1a+/- mice. We first demonstrated that eEF2K deletion per se did not cause any behavioral alteration. Our behavioral characterization demonstrated that eEF2K-/- mice behave like WT mice in motor coordinator tests and in cortex-, hippocampus- and amygdala-dependent behavioral paradigms presenting only a mild hippocampal-dependent phenotype in the context of fear conditioning (Heise et al., 2017). Thus, we confirmed that Scn1a+/- mice recapitulated the majority of comorbidity found in Dravet syndrome patients. In particular, they showed impairments in motor coordination, with no alteration in walking and climbing or in muscle strength. Moreover Scn1a+/- mice compared to WT mice had a reduced memory for objects and their spatial relationships. These memories and learning problems were also confirmed by the T-maze and Morris water maze test where Scn1a+/- mice performed worse than WT mice during the reversal phase of the test. Scn1a+/- mice also presented stereotyped behaviors although they did not show anxiety-like or major depression-like behaviors. Interestingly, all these behavioral alterations were rescued by the deletion of the eEF2K gene strongly suggesting that the inhibition of the eEF2K/eEF2 pathway by increasing the strength of the GABAergic transmission was sufficient not only to rescue the epileptic phenotype of Scn1a+/- mice but also the associated comorbidity.

These results and our previous study, demonstrating that genetic inhibition of eEF2K rescued the epileptic phenotype in a different model of genetic epilepsy (Heise et al., 2017), indicate eEF2K as a possible target for develop new pharmacological the treatment for epilepsy. Thus, we tested the effect of pharmacological inhibition of eEF2K on Scn1a+/- mice. Our results demonstrat that subcronic treatment with A484954, a selective eEF2K inhibitor, was able to ameliorate the epileptic phenotype of the Scn1a+/- mice by reducing eEF2 phosphorylation in the hippocampus and cerebral cortex. Even though future studies focused on clarifying the activity of eEF2K inhibition on Dravet syndrome comorbidity are needed, our data suggest eEF2K pharmacological inhibition as an innovative therapy for Dravet syndrome.

In conclusion, Scn1a+/- mice exhibited increased levels of eEF2 phosphorylation that were associated with EEG alterations and a greater responsiveness to seizures accompanied by behavioral abnormalities including memory impairment, ASD like behavior and motor coordination problems. eEF2K deletion in Scn1a+/- mice was sufficient to rescue all these impairments enhancing GABAergic transmission and modulating the PI3K/Akt/mTOR pathway. Despite the fact that eEF2k is an ubiquitous and conserved protein it has only one substrate (Liu and Proud, 2016) and its deletion in WT mice caused only minor impairments suggesting that inhibiting its function should not elicit serious side effects. Notably pharmacological inhibition of eEF2K in adult mice was also sufficient to normalize the EEG profile of Scn1a+/- mice proposing eEF2K as a new pharmacological target not only for Dravet syndrome but also for a number of neuronal diseases in which eEF2K activity is altered (Beretta et al., 2020).

## Material and Methods

### Mice

*Scn1a*^+/-^ *eEF2K*^-/-^ mice were generated by crossing *Scn1a*^+/-^ mice (Yu et al., 2006) with *eEF2K*^-/-^ mice (Ryazanov, 2002) and backcrossed for about 20 generations before were used for all the experiments. Mice were housed under constant temperature (22 ± 1°C) and humidity (50%) conditions with a 12 h light/dark cycle and were provided with food and water ad libitum. All the experiments were performed on female and male mice. All experiments involving animals followed protocols in accordance with the guidelines established by the European Communitis Council and the Italian Ministry of Health (Rome, Italy) for the correct use of laboratory animals in research. Experimental procedures of EEG and behavioral analysis followed the guidelines established by the Italian Council on Animal Care and were approved by the Italian Government decree No. 17/2013 and 980/2017. All efforts were made to minimize the number of subjects used and their suffering.

#### Mice genotyping

All primers were provided from Thermo Fisher and the REDExtract-N-Amp PCR Reaction Mix™ reagent used for the polymerase reaction was provided from Sigma-Aldrich^®^. The PCR for eEF2K was performed using two pairs of primers SA8 (5’-GGCCGGCTGCTAGAGAGTGTC-3’) and SA5 (5’-CATCAGCTGATTGTAGTGGACATC-3’) for WT allele and SA8 and Neo1 (5’-TGCGAGGCCAGAGGCCACTTGTGTAGC-3’) for mutated allele. PCR for SCN1A was performed using the following primer: 268 (5’-CGAATCCAGATGGAAGAGCGGTTCATGGCT-3’), 269 (5’-ACAAGCTGCTATGGACATTGTCAGGTCAGT-3’) and 316 (5’-TGGCGCTCGATGTTCAGCCCAAGC-3’).

### Protein biochemistry

Mice were sacrified and hippocampus, cerebral cortex, liver, kidney and heart were dissected and lysate in buffer containing 10 mM Hepes pH 7.4, 2 mM EDTA, protease inhibitors (Sigma, P8340) and phosphatase inhibitors (Roche). Samples were centrifuged at 800 x g for 5 min at 4°C. Resulting supernatant were collected and quantified by BCA protein assay (EuroClone) to assess protein concentration and then solubilized in 4x loading dye (250 mM Tris-HCl pH 6.8, 40% glycerol, 0.008% bromophenol blue, 8% SDS; all from Sigma-Aldrich). All samples were boiled at 65°C for 10 minutes and then equal amounts of each sample were separated using SDS-PAGE and subsequently blotted on nitrocellulose membranes using the Trans-Blot Turbo System (Bio-Rad). Membranes were washed in Tris-buffered saline-Tween (TBS-T) (20 mM Tris pH 7.4, 150 mM NaCl (both Sigma-Aldrich) and 0.1% Tween 20 (Bio-Rad). After 1 hour blocking at room temperature with 5% Bovine Serum Albumin (BSA) or 5% milk in TBS-T, membranes were incubated overnight at 4°C with primary antibody in blocking buffer (TBS-T containing 3% BSA or 3% milk). The membranes were washed three times in TBS-T and then incubated with HRP-conjugated secondary antibodies in TBS-T and 3% BSA or 3% milk for 1 hour at room temperature. After three washes (10 min each), chemiluminescence was induced using an ECL Western Blotting Substrate kit and further detected using a ChemiDoc XRS+ machine. All signals were quantified using ImageLab softwer and normalized against the values of the respective signal for actin or βIII-tubulin.

### Antibodies

The following primary antibody were used (dilution and source): mouse anti-actin (1:5.000 Sigma-Aldrich), mouse anti-βIII-tubulin (1:10.000 Sigma-Aldrich), rabbit anti-eEF2 (1:500 Cell Signaling), rabbit anti-peEF2 (T56) (1:1.000 Cell Signaling), mouse anti-Akt (1:1.000 Cell Signaling), rabbit anti-pAkt (S473) (1:1.000 Cell Signaling), rabbit anti-synapsin1/2 (1:1000 Synaptic Systems), mouse anti-GABA_A_R ***α***5 (1:250 NeuroMab). All HRP-conjugated secondary antibodies were purchased from Jackson ImmunoResearch. HRP anti-rabbit (1:3.000) and anti-mouse (1:5.000) were used for western blot.

### Electrophysiology

#### Preparation of brain slices

Hippocampal slices were prepared from P25–P35 mice using standard procedures (Liautard et al., 2013; Zerimech et al., 2020). Mice were deeply anesthetized with isoflurane and decapitated. The brain was quickly removed, and horizontal hippocampal slices (300 μm) were cut with a Vibratome in chilled (0–4°C) slicing solution containing (mM) 75 sucrose, 87 NaCl, 25 NaHCO_3_, 25 D-glucose, 2.5 KCl, 1.25 NaH_2_PO_4_, 0.5 CaCl_2_, 7.0 MgCl_2_. The slices were transferred to a storage chamber andincubated at room temperature for at least 45 min before recording, then transferred to a recording chamber and perfused with ACSF containing (mM) 129 NaCl, 3 KCl, 1.8 MgSO_4_, 1.6 CaCl_2_, 1.25 NaH_2_PO_4_, 21 NaHCO_3_, and 10 D-glucose at 28/30 °C. All solutions were saturated with 95% O_2_ and 5% CO_2_.

#### Brain slice recording

Whole-cell voltage-clamp recordings were performed on CA1 hippocampal pyramidal neurons visualized with near infrared differential interference contrast (DIC) optics.For the recordings of inhibitory postsynaptic currents (IPSCs), the intracellular solution contained (mM) 135 CsCl, 10 N-2-hydroxyethylpiperazine-N′-2-ethanesulfonic acid (HEPES), 0.2 ethylene glycol-bis(β- aminoethylether)-N,N,N′,N′-tetraacetic acid (EGTA), 2 Mg-ATP, 4 ATP, 10 phosphocreatine, pH 7.2 with CsOH. Kynurenic acid (KA, 3 mM) was included in the extracellular recording solution to block the excitatory synaptic transmission in IPSC recordings. The holding potential was -70 mV. Patch pipettes were pulled from borosilicate glass and, when filled with the intracellular solution, their resistance ranged from 3 to 5 MΩ. Recordings were obtained by means of a Multiclamp 700B amplifier, Digidata 1440 digitized and pCLAMP 8.0 software (Molecular Devices). Access resistance was continuously monitored for each cell. Only the cells with access resistance less than 20 MΩ were recorded, and recordings were terminated/discarded when a significant (>10%) increase occurred. Data were analyzed using Clampfit 10.5 (Molecular Devices) and Mini Analysis (Synaptosot).

### Electroencephalography analysis

For electroencephalography (EEG) analysis P60-P90 mice were used. Mice were anesthetized with isoflurane (2% (v /v) in 1 L/min O2). Four screw electrodes (Bilaney Consultants GMBH, Dusseldorf, Germany) were inserted bilaterally through the skull over the cortex (anteroposterior, +2.0 –3.0 mm; let–right 2.0 mm from bregma), as previously described (Manfredi et al., 2009) and in agreement to brain atlas coordinates (Franklin and Paxinos, 2008).;Mice were allowed to recover for approximately a week from surgery under antimicrobial cover (cetriaxone, Sigma-Aldrich; 50 mg/kg i.p.) before performing the experiments. EEG activity was recorded in awake freely moving mice placed in a Faraday chamber using a Power-Lab digital acquisition system (AD Instruments, Bella Vista, Australia; sampling rate 100 Hz, resolution 0.2 Hz).

#### Electroencephalography analysis under baseline conditions

Seven days after surgery basal cerebral activity was recorded continuously for 24 hours. All EEG tracings were analyzed and scored for the presence of spikes. EEG spikes were recognized as having a duration <200 ms with a baseline amplitude set to 4.5 times the standard deviation of the EEG signal (determined during inter-spike activity periods, whereas repetitive spiking activity was defined as 3 or more spikes lasting <5 seconds). Segments with electrical noise or movements artifacts were excluded from statistical analysis (Manfredi et al., 2009).

#### Electroencephalography analysis under thermal stress condition

The day afterEEG basal activity, mice body temperature was monitored and managed by a rectal temperature probe connected to a controller. Average mouse body temperature is 36.9°C and it was controlled by a heat lamp above the chamber. Mice were accustomed to the Plexiglas cage for at least 10 minutes at 37.5°C. Then, the body temperature was increased by 0.5°C every 2 minutes until a tonic-clonic seizure occurred or 42.5°C was reached (Oakley et al., 2009; Salgueiro-Pereira et al., 2019). Immediately after the heat stimulus was removed. Seven days after heat EEG basal activity was measured for 24 h and the number of spikes was evaluated.

#### Electroencephalography analysis under pharmacological inhibition of eEF2K by A484954

Two days after monitoring basal cerebral activity for 24 hours pellets were implanted in the lateral side of Scn1a+/- mice between ear and shoulder. Animals were treated for ten days with 10 mg of A484954/pellets that release 1 mg of drug every day. Placebo pellets were used in experiment as the proper control. During the 10 days of treatment EEG activity was recorded for 24 hours after 3,6 and 9 days of pellets implantation and analysed as previously described. Animals were sacrificed the last day of the pharmacological treatment.

### Behavioral procedure

Mice of P60-P90 were used for behavioral experiments. Animals were housed in groups of four or five individuals of mixed genotypes. All of the tests were conducted during the light portion of the cycle. To minimize the number of animals each mouse was submitted to a maximum of 3 tests with an interval of 1 week between two tests. All the behavioural experiments followed the ARRIVE guidelines. A total of 205 mice was used (64 WT,76 Scn1a+/-,65 Scna1+/-eEF2K-/-. Animals were divided in different groups (Group 1 – EEG 4WT,6 Scn1a+/-,5Scna1+/-eEF2K-/- for heat seizure, 5 Scn1a+/- for vehicle, 5 Scn1a+/- for A484954); Group 2 -behavioural tests, 10 mice per genotype: 2A-elevated plus maze, novel object recognition, tail suspension ; 2B-self-grooming,wire hanging, spatial object recognition),Group 2C (balance beam, marble burying, pole test);Group 2D (locomotor activity, sociability/social novelty, rotarod) Group 2E: Morris water maze, acquisition and reversal; Group 2F (T-maze, acquisition and reversal).

In addition, a further group of 70 animals (34 WT and 36 eEF2K-/-) was used to investigate sensory abilities, motor coordination, sociability, anxiety- and depression-like behaviour and episodic and spatial memory (Group3A (range of 8-12 animals for genotype): olfactory and visual cliff test, balance beam and Novel Object Recognition; Group 3B (a range of 5-10 animals for genotype): Self-grooming, Pole test and Spatial Object Recognition; Group 3C (range of 5-12 animals for genotype): Rotarod test, Sociability test and Tail Suspension). All the behavioral scoring was performed on a blind basis by a trained experimenter.

#### Locomotor activity

The spontaneous motor activity was evaluated in an automated activity cage (43 × 43 × 32 cm) (Ugo Basile, Varese, Italy), located in a sound-attenuating room as previously described (Braida and Sala, 2000). Horizontal and vertical activity were detected by infrared photobeam break sensors put from 2.5 cm and 4 cm from the floor of the cage, respectively. Cumulative horizontal and vertical movements were counted for 3 hours.

#### Wire hanging

The limb force was tested by positioning a mouse on the top of a wire cage lid (19 × 29 cm) that was turned upside down at approximately 25 cm above the surface of the bedding material. The grip of the mouse was ensured by gently waving three times before rotating the lid as described in (Oliván et al., 2015). The latency to fall onto the bedding was registered (cut-off = 300 s).

#### Balance beam walking test

The balance beam walking test is a test for assess motor coordination and balance as previously described (Luong et al., 2011). The beam apparatus consisted of 1 m beams with a flat surface of 12 mm width resting 50 cm above the table top on two poles. At the end of the beam a black box was placed as the finish point. Nesting material from home cages was placed in the black box to attract the mouse to the finish point. A lamp (with 60-watt light bulb) was used to shine light above the start point and served as an aversive stimulus. A video camera was set on a tripod to record the performance. On training days, each mouse crossed the 12 mm beam 3 times. The time required to cross to the escape box at the other end (80 cm away) was measured with a stopwatch. The stopwatch started when the nose of the mouse began to cross the beam and stopped when the animal reaches the escape box. Once the mice are in the safe box, they are allowed some time (∼15 sec) to rest there. Before the next trial the mice rest for 10 min in their home cages between training sessions on the two beams. On the test day, times to cross each beam were recorded. Two successful trials in which the mouse did not stall on the beam are averaged. The beams and box were cleaned of mouse droppings and wiped with towels soaked with 70% ethanol and then water before the next beam was placed on the apparatus.

#### Pole test

In the pole test, which evaluated motor coordination, mice were trained for 2 days (in the morning and in the afternoon) to descend a vertical pole (90 cm length, 1 cm diameter). Every training consisted of three trials. Mice were accustomed to the room 20 minutes before trials and test day. The test was performed the third day and the time necessary to the mice to descend the pole in five trials was recorded. A cut-off of 60 seconds was given (Rial et al., 2014), with minor modifications). Data were shown as mean of 5 trials evaluated during the test day.

#### Rotarod

The rotarod was used to measure fore and hindlimb motor coordination and balance. During the training period, each mouse was placed on the rotarod apparatus (Ugo Basile, Biological Research Apparatus, Varese, Italy) at a constant speed 12 rpm for a maximum of 120 seconds, and the latency to fall off the rotarod within this time period was recorded. Mice received four trials per day for 4 straight days.

#### Repetitive self-grooming

Mice were assessed for spontaneous self-grooming as a measure of repetitive behavior as previously described in (McFarlane et al., 2008). Each mouse was placed into a standard plastic cylinder (46 × 23.5 × 20 cm). Cylinder was empty to eliminate digging in the bedding, which is a potentially competing behavior. The room was illuminated at about 40 lux. A front-mounted CC TV camera (Security Cameras Direct) was placed at circa 1 m from the cages to record the sessions. Sessions were video-taped for 20 min. The first 10 min of habituation was not scored. In the next 10 minutes mice were scored with a stopwatch for cumulative time spent grooming all the body regions. The cylinder was cleaned with 70% ethanol between each animal.

#### Elevated plus maze

The Elevated Plus Maze paradigm was used to study anxiety related behavior. The apparatus had a central platform (10 × 10 cm) from which originated two opposite open arms (30 × 10 cm) and two enclosed arms (30 × 10 × 14 cm) according to (Lister, 1987). The apparatus was made of white wood, placed to a height of 60 cm above floor level in the center of a small quiet room under dim light (about 30 lux). The test was conducted in the morning, during the early light phase of the light cycle. After 20 minutes of familiarization to the novel environment, mice were placed individually onto the center of the apparatus, facing an open arm. Experimenter recorded for 5 minutes the number of open- and closed-arm entries and the time spent in open- and closed-arms.

#### Marble burying

Mouse marble burying can be associated to both anxiety-like traits, increased by novelty, and obsessive/compulsive-like behaviour as previously described in literature (Deacon, 2006). The marble burying assay is applied to evaluate how many novels glass marbles a rodent would bury. This behaviour was assessed using a clean cage (50 × 25 × 30 cm) filled with 3 cm bedding, lightly tamped down to make a flat, even surface. Each mouse was placed individually into the corner of the empty cage where previously food marbles were placed covered by the bedding. The latency to the first marble buried and the number of marbles buried (to 2/3 their depth) with bedding were recorded over a maximum period of 900 sec.

#### Tail suspension

Tail suspension test assayed depressive-like behaviors of mice as described in (Steru et al., 1985). Tail suspension test was performed on an apparatus with a support at 35 cm from the basis where was fixed a hook where mouse tail was fasten at about 1 cm from the origin. The test was preceded by a familiarization phase where mice were let in the room test at least 1 hour before. The test day lasted 6 min during which the experimenter measured time of mouse immobility.

#### Novel Object Recognition test

The novel-object recognition test was performed over 3 straight days in an open plastic arena (60 × 50 × 30 cm) as previously described (Pan and Xia, 2008). The test had 3 phases. In the habituation one, the first day, mice were habituated to the empty arena for 10 minutes, the familiarization and novel-object recognition the day after. In the familiarization phase, two identical objects were placed in the middle of the arena equidistant from the walls and from each other. Mice were placed between the two objects until it had completed 30 s of cumulative object exploration (20 minutes cut-off). Object recognition was measured when each animal was within approximately 1 cm of an object with its nose toward the object. Climbing the object or pointing the nose toward ceiling near the object were not considered exploring behaviors. After familiarization, mice were returned to the home cage until they were tested for novel recognition after 5 minutes, 120 minutes or 24 h. In the novel recognition phase, a novel object (never seen before) took the place of one of the two familiars. Scoring of object recognition was performed as during the familiarization phase. For each mouse, the role (familiar or new object) as well as the relative position of the two objects were randomly permuted. The objects used for the test was white plastic cylinders and colored plastic Lego stacks of different shapes. The arena was cleaned with 70% ethanol after each trial. Performance was analyzed by calculating a discrimination index (N-F/N+F), where N = the time spent exploring the novel object, and F = the time spent exploring the familiar object.

#### Spatial object recognition test

Spatial object recognition test was performed in an arena according to (Kenney et al., 2011), with minor modifications that consisted in an opaque white plexiglass cage (58 × 50 × 43 cm) that was dimly lit from above (27 lux) and two visual cues were placed above two adjacent walls. In the center of the northern wall there was a black and white stripped pattern (21 × 19.5 cm) and in the center of the western wall there was a black and gray checkered pattern (26.5 × 20 cm). The objects were placed across the visual cues. Mice were habituated to the arena for 10 minutes the day before the test. The test day, first mice were allowed to familiarize with two different objects. The experimenter measured the time spent in sniffing both objects until the mouse completed 30 seconds in exploring objects (cut-off 20 minutes). Exploring behavior was defined as mouse having its nose directed toward the object and within approximately 1 cm of the object (Bevins and Besheer, 2006); climbing or sitting were not considered exploration behaviors. After 5 minutes, 120 minutes and 24 hours from familiarization phase mice were allowed to re-explore the cage where one object was moved in a new position. Scoring of object recognition was performed as during the familiarization phase. Between two sessions, mice returned to their home-cage. Cage and object were carefully cleaned with 70% ethanol before and after all behavioral procedures. Performance was analyzed by calculating a discrimination index (N-F/N+F), where N = the time spent exploring the moved object during the test and F = the time spent exploring the unmoved object during the test.

#### T-maze

Mice were deprived of food until they reached 90% of their free-feeding body weight. Mice were habituated to a black wooden T-maze (with a 41 cm stem section and a 91 cm arms section, and each section was 11 cm wide and had walls that were 19 cm high) and trained to obtain food within the maze for 5 days as previously described (Sala et al., 2011). During the acquisition phase, one arm was designated to be reinforced with Coco Pops (Kellogg’s) in each of ten daily trials. Each mouse was placed at the start of the maze and allowed to freely move and choose which arm to enter. The number of days required to reach the goal criterion (80% correct for 3 days) was recorded. Each mouse that met the goal for acquisition was then tested using a reversal procedure in which the reinforce was switched to the opposite arm. A cut off of 20 days in both acquisition and reversal phase was established.

#### Morris water maze

The Morris water maze test was used to analyze changes in the learning and memory abilities of the mice according to the methods described in Morris (Morris, 1984) (adapted for mice). The Morris water maze consisted of a circular water maze (120 cm in diameter x 50 cm in height) filled with water. The pool was divided into 4 quadrants. A circular hidden platform with a diameter of 10 cm was placed inside the maze, and its surface was maintained at 0.5 cm below the surface of the water.

Floating plastic particles were placed on the surface of the water to hide the platform from sight according to the methods of (Zhang et al., 2013). During the habituation trials, mice were placed in a random area inside the maze and allowed to swim for 60 seconds. During the acquisition trials, mice were given 4 daily trials (with 60 minutes inter-trial intervals) for 4 straight days in which each mouse empty wire cage was released into the pool at different starting points and trained to locate a constant platform position. At 24 hours after the last trial, a probe test was performed in which the platform was removed. Two days later, a reversal task was performed to assess cognitive flexibility. The platform was placed in the opposite quadrant of the maze, and 4 daily trials were performed for 4 straight days. On the fifth day, a probe trial was performed that was similar to that in the acquisition phase. The time spent in the target area and the latency to reaching the target zone were evaluated.

#### Sociability and preference for social novelty test

The sociability tests were performed in a rectangular apparatus in transparent polycarbonate with three-chamber (width = 42.5 cm, height = 22.2 cm, central chamber length = 17.8 cm, and side chamber lengths = 19.1 cm) as previously described in (Sala et al., 2011). In the 10 minutes habituation phase, each mouse was placed in the middle compartment and free to explore all chambers. Then, one side of the compartment was occupied an unfamiliar male mouse and the other contained an empty wire cage. Immediately after sociability test, without cleaning the apparatus, it was performed the social novelty test putting an unfamiliar mouse in the empty wire cage. Every test lasted 10 minutes in which were measured the time spent exploring each chamber. The data were expressed in sociability index (SI) and social novelty preference index (SNI) as follows: SI = (time exploring novel mouse 1 – time exploring empty cage)/(time exploring novel mouse 1 + time exploring empty cage) and SNI = (time exploring novel mouse 2 – time exploring familiar mouse)/(time exploring novel mouse 2 + time exploring familiar mouse).

### Statistical analyses

Based on the number of comparisons and the pattern of data distribution, appropriate statistical tests were used to analyze the data. Data are expressed as the mean ± SEM or percentage, analyzed for statistical significance, and displayed by Prism 8 software (GraphPad Software). Shapiro–Wilk tests was applied to test the normal distribution of experimental data. Normal distributions were compared with one simple t-test, student t-test or ANOVA with appropriate post hoc test. Non-normal distributions were compared with the non-parametric Wilcoxon test, Mann–Whitney test or Kruskal – Wallis test with appropriate post hoc test, as indicated. The accepted level of significance was p<0.05. Statistical analyses for every experiment are described accordingly in the figure legends.

## Acknowledgement

This work was supported by Fondation Jerome Lejeune (project #1638) to C.S. the Comitato Telethon Fondazione Onlus (grant no. GGP16131 to C.V. and GGP17176 to C.S) and Regione Lombardia NeOn Progetto “NeOn” POR-FESR 2014-2020, “, ID 239047, CUP E47F17000000009 to C.S and C.V.

## Author contributions

Stefania Beretta, Luisa Ponzoni, Laura Gritti and Paolo Scalmani performer all the experiments; Stefania Beretta and Luisa Ponzoni prepared the figures; Massimo Mantegazza supervised the electrophysiology experiments, Mariaelvina Sala supervised all the behavioral experiments; Chiara Verpelli and Carlo Sala supervised and decided the experiments, wrote the paper; all authors contributed to edit the text and figures.

## Figure legends

**Supplementary Figure 1.**
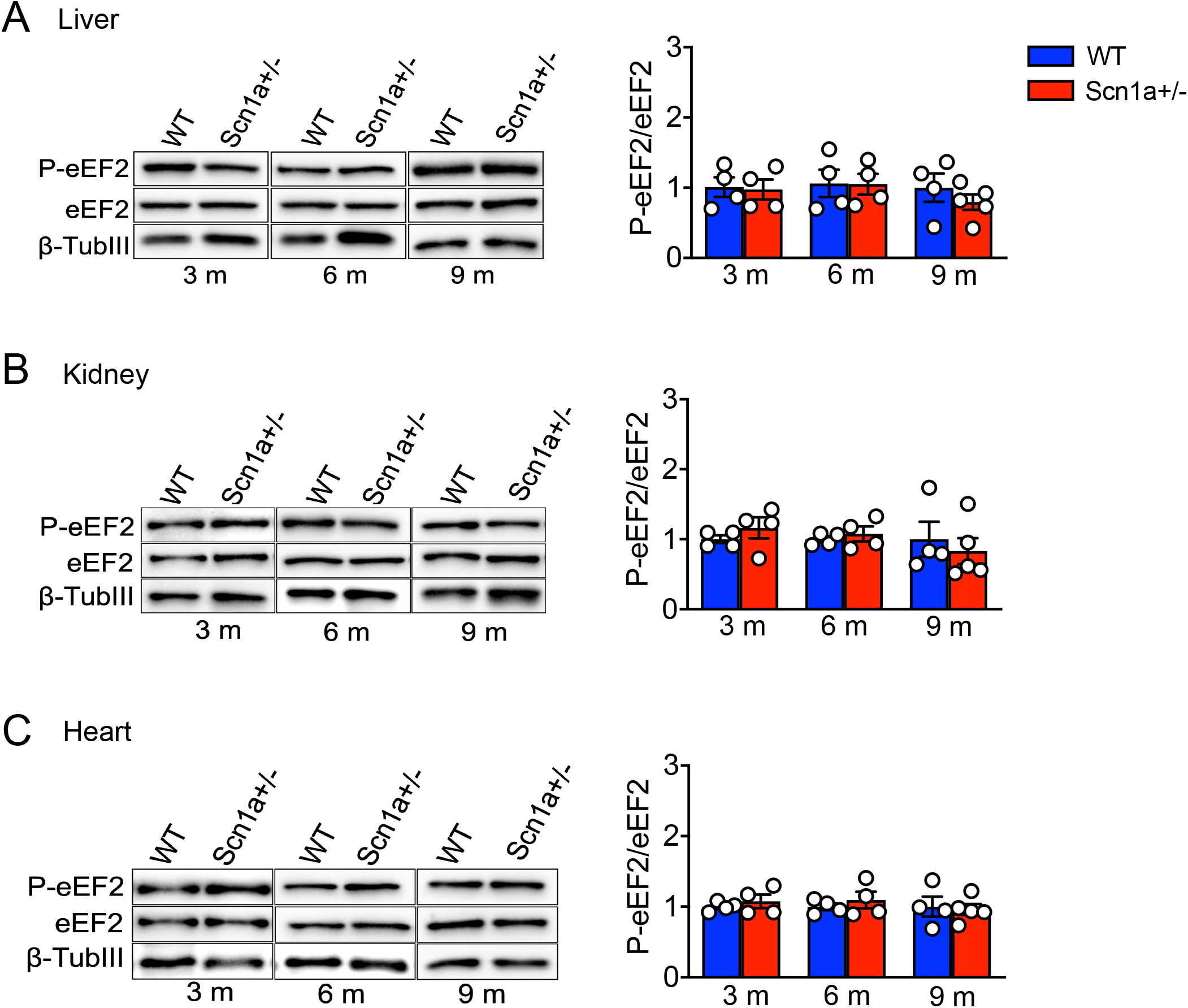
No difference in eEF2 phosphorylation levels in total homogenate of liver, kidney, and heart between Scn1a+/- and WT mice. **(A-C)** Western blots and relative quantification for phosphorylated eEF2 in samples of liver **(A)**, kidney **(B)** and heart **(C)** show no significant difference in 3, 6 and 9 months old in Scn1a+/- mice compared with WT mice. All Data are presented as mean ± SEM; N=4 per group. One-sample *t*-test.

**Supplementary Figure 2.**
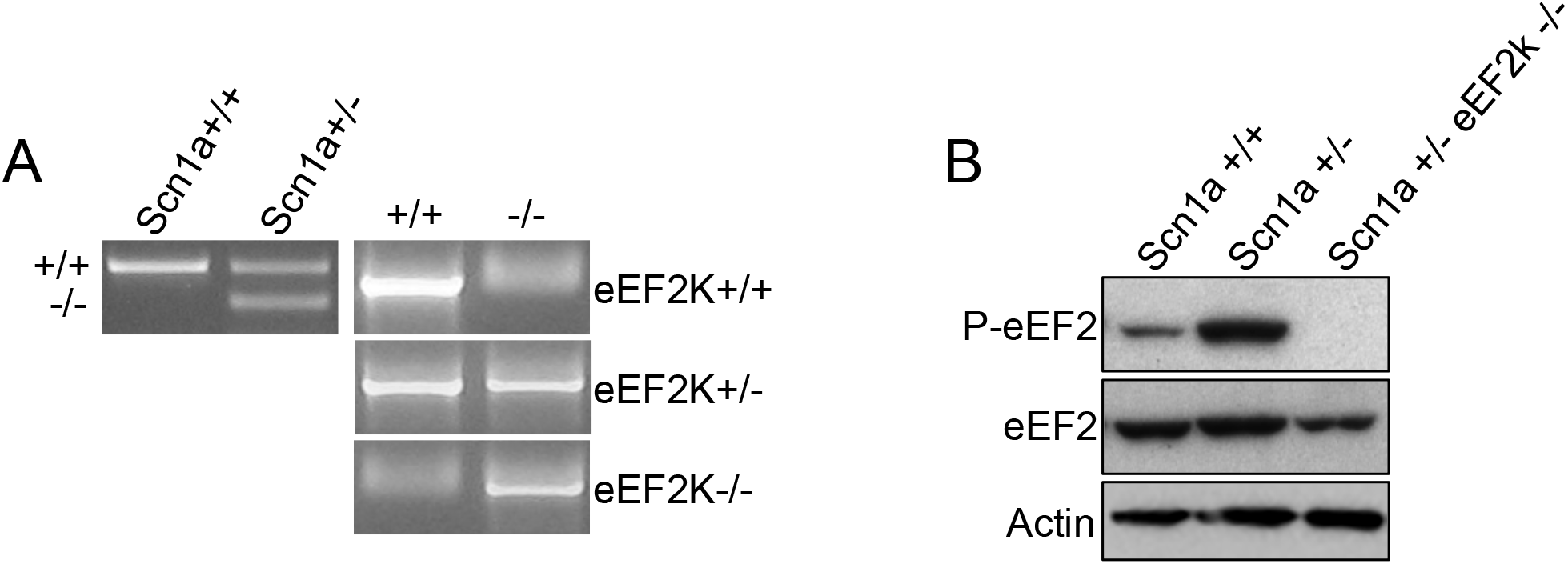
Genetic characterization and levels of phosphorylated eEF2 in WT, Scn1a+/- and Scn1a+/-eEF2K-/- mice. **(A)** Representative PCR for SCN1A gene (left). WT Scn1a+/+ (WT) mice display single band of 300 bp, heterozygous Scn1a+/- mice display one at 300 bp and the other ad 150 bp. Representative PCR for eEF2K gene (right). Length of the bands for WT and KO is at the same high at 1.2 kbp. Two different PCR were performed for WT and KO. **(B)** Western blot for phosphorylated eEF2 in WT (Scn1a+/+), Scn1a+/- and Scn1a+/-eEF2K-/- shows that eEF2 phosphorylation is totally absent in Scn1a+/-eEF2K-/- mice.

**Supplementary figure 3.**
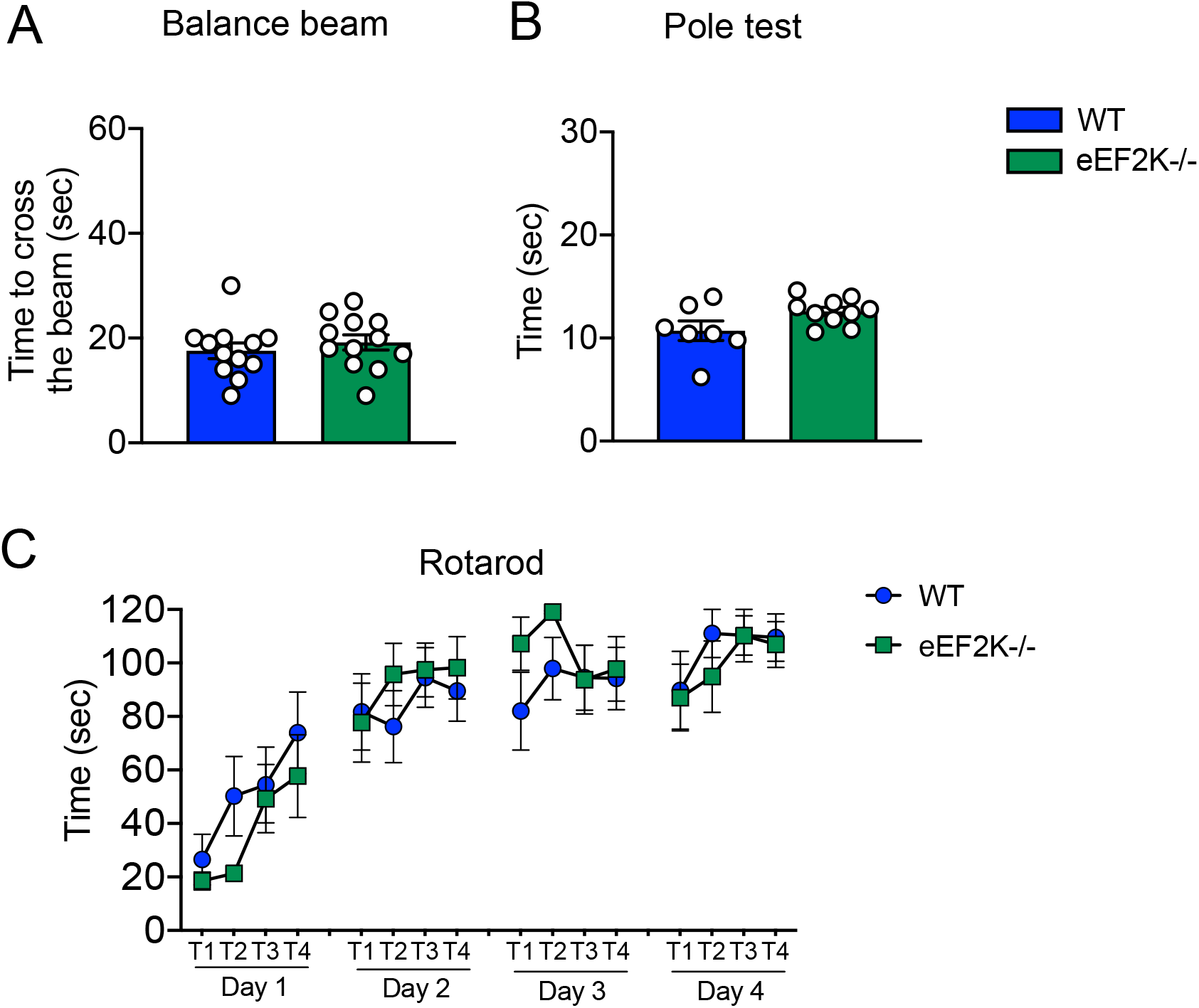
Genetic deletion of eEF2K does not alter balance and motor coordination in WT mice. **(A)** eEF2K-/- mice show no impaired in balance compared with WT when cross the 12 mm beam in balance beam test. Data are presented as mean ± SEM. N=12 per group. Unpaired two-tailed *t*-test. **(B)** eEF2K-/- mice behave like WT in pole test measured by the time take to downstream the pole. Data are presented as mean ± SEM. WT n=7, eEF2K-/- n=10. Unpaired two-tailed *t*-test. **(C)** eEF2K-/- display normal motor coordination when tested by rotarod test. Data are presented as mean ± SEM. N=10 per group. Unpaired two-tailed *t*-test and two-tailed Mann-Whitney test.

**Supplementary Figure 4.**
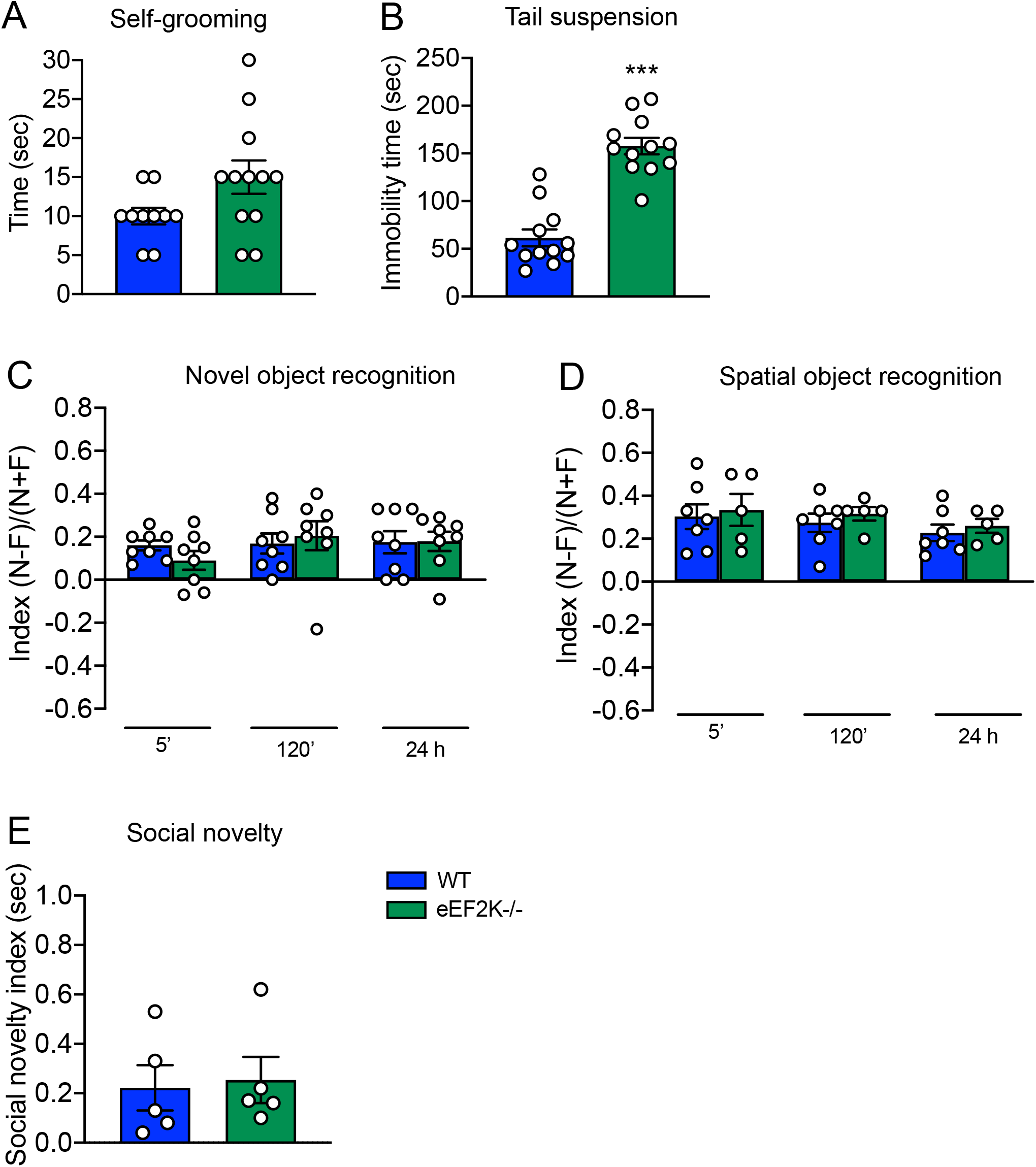
eEF2K-/- mice show depressed-like behaviour but no other autistic behaviours when compared with WT. **(A)** No differences in repetitive grooming behavior evaluated in terms of time spent to do grooming between WT and eEF2K-/- mice. Data are presented as mean ± SEM. WT n=10, eEF2K-/- n=12. Unpaired two-tailed *t*-test with Welch’s correction. **(B)** eEF2K-/- mice show depressed-like behavior in tail suspension test measured by the time spent immobile. Data are presented as mean ± SEM. N=12 per group. Statistical analysis ***p<0.001; Unpaired two-tailed *t*-test. **(C-D)** eEF2K-/- mice do not show alteration in recognition memory **(C)** and spatial memory **(D)** evaluated by the discrimination index in the novel object recognition test **(C)** and spatial object recognition test **(D)** at 5, 120 min and 24h after familiarization phase. Data are presented as mean ± SEM. N=8 per group in novel object recognition test. WT n=7, eEF2K-/- n=5 in spatial object recognition. Unpaired two-tailed *t*-test; Two-tailed Mann-Whitney test (120 min in novel object recognition test). **(E)** eEF2K-/- behave as WT mice in social interaction. Social novelty preference index (right). All data are presented as mean ± SEM. N=5 per group; Two-tailed Mann-Whitney test.

**Supplementary figure 5.**
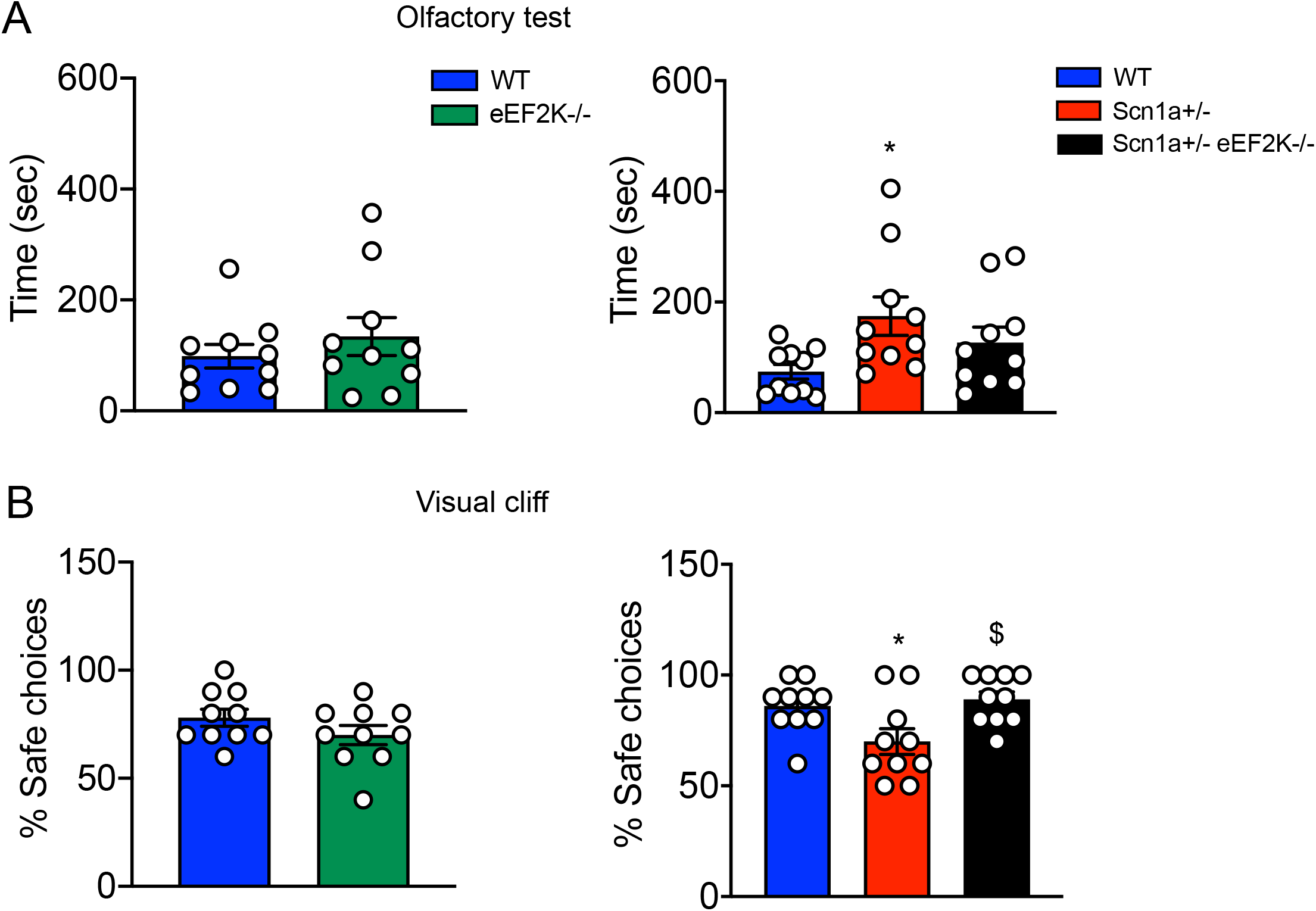
eEF2k deletion improve olfactory ability and visual acuity of Scn1a+/- mice. **(A)** Shows the comparison of the time spending to find the food between WT and eEF2K-/- (left) and between WT, Scn1a+/- and Scn1a+/- eEF2K-/- (right). Similar behaviour between WT and eEF2K-/- mice (left). Data are presented as mean ± SEM. N=10 per group; Unpaired two-tailed *t*-test. Scn1a+/- mice spent more time to find the food when compared with WT mice (right). Data are presented as mean ± SEM. N=10 per group. Statistical analysis *p<0.05; One-way ANOVA, Tukey’s post hoc. **(B)** Shows the comparison of the % of safe choices between WT and eEF2K-/- (left) and between WT, Scn1a+/- and Scn1a+/- eEF2K-/- (right). Similar visual acuity for WT and eEF2K-/- mice (left). Data are presented as mean ± SEM. N=10 per group; Unpaired two-tailed *t*-test. Scn1a+/- show a poor visual acuity when compared with WT and Scn1a+/-eEF2K-/- mice (right). Data are presented as mean ± SEM. N=10 per group. Statistical analysis *p<0.05 versus corresponding WT, $p<0.05 versus corresponding Scn1a+/-; One-way ANOVA, Tukey’s post hoc.

